# Bumped kinase inhibitor BKI-1708 interferes in cytokinesis and drives baryzoite stage conversion in the cyst-forming apicomplexan parasites *Toxoplasma gondii, Neospora caninum* and *Besnoitia besnoiti*

**DOI:** 10.1101/2025.10.22.684074

**Authors:** Maria Cristina Sousa, Joachim Müller, Kai Pascal Alexander Hänggeli, Manfred Heller, Anne-Christine Uldry, Sophie Braga-Lagache, Alexandre Leitao, Luis-Miguel Ortega-Mora, Kayode K. Ojo, Wesley C. Van Voorhis, Andrew Hemphill

## Abstract

Bumped kinase inhibitors (BKIs) have demonstrated safety and promising efficacy against various apicomplexan pathogens both *in vitro* and *in vivo*. However, in the closely related cyst-forming coccidians *T. gondii*, *Neospora caninum* and *Besnoitia besnoiti*, *in vitro* treatments with a range of BKIs induced the conversion of intracellular tachyzoites into atypical multinucleated complexes (MNCs), also named “baryzoites”. In this study, baryzoites of *T. gondii, N. caninum* and *B. besnoiti* generated through exposure of tachyzoites to 2.5 µM BKI-1708 were comparatively assessed. TEM showed that baryzoites contained multiple nuclei, clustered together and separated from the cytoplasmic organelles of newly formed zoites. These zoites do not have outer tachyzoite plasma membrane, were unable to complete cytokinesis, remained intracellular, and were enclosed by a parasitophorous vacuole membrane. TEM demonstrated the presence of an electron-dense cyst wall-like components only in *T. gondii* baryzoites. Species-specific differences in antigen expression were observed by immunofluorescence using specific antibodies. Comparative proteomic analysis revealed consistent downregulation of ribosomal proteins, proteins associated with secretory organelles, as well as of transcription and translation factors in all baryzoites. Bradyzoite-specific markers were upregulated only in *T. gondii* baryzoites. In addition, common orthologues of two alveolin-domain filament proteins (IMC7 and IMC12) and a hypothetical protein (TGME49_236950, NCLIV_050850, BESB_060040) were detected at higher abundance in all treated parasites. Overall, baryzoites exhibit distinct phenotypic and proteomic profiles, with ambiguous expression of tachyzoite and bradyzoite antigens, and lacking complete cellular division under drug pressure, suggesting a reversible response to stress rather than progression into a fully differentiated form.

**Significance:** Apicomplexan parasites cause serious diseases worldwide, yet treatment options remain limited. A promising group of drugs are BKIs. We investigated how BKI-1708 affects threclosely related *T. gondii*, *N. caninum*, and *B. besnoiti*. Instead of killing the parasites, the drug induced the formation of multinucleated structures termed “baryzoites”. These baryzoites exhibited ambiguous characteristics during the actively growing and dormant stages of the parasite life cycle and were unable to complete normal cell division. Moreover, we observed other key similarities and differences among species including downregulation of ribosomal proteins and transcription/translation factors, while only *T. gondii* displayed cyst wall formation. Microscopy and proteomics demonstrated that baryzoites represent a distinct stage that is formed upon drug pressure and promotes parasite survival during prolonged drug exposure. These findings highlight the unexpected ways parasites adapt to drug treatment and provide new insights into how BKIs exert their activities.

## Introduction

The subphylum Apicomplexa (phylum Alveolate) includes several intracellular parasites of significant medical and veterinary importance, among which the cyst-forming coccidia *Toxoplasma gondii*, *Neospora caninum*, and *Besnoitia besnoiti* form a closely related cluster within the family Sarcocystidae (1). The genomes of *T. gondii, N. caninum* and *B. besnoiti* exhibit a high degree of synteny and encode a large number of known and inferred orthologues that are important for critical processes such as host cell invasion, immune modulation, intracellular survival as well as sexual development (2–4).

*T. gondii* and *N. caninum* undergo a life cycle that alternates between intermediate and definitive hosts. The intermediate hosts harbor two stages: (i) the rapidly proliferating and disease-causing tachyzoites, which undergo continuous lytic cycles and thus inflict tissue damages and immunopathology, and (ii) the slowly proliferating bradyzoites that form long-lived tissue cysts mostly in the central nervous system and muscle tissues, representing the chronic stage of infection which normally does not cause clinical symptoms. Within the intestine of the definitive hosts - felids in the case of *T. gondii* and canids for *N. caninum* - sexual development results in the production of oocysts that are shed with the feces, and sporulation in the environment leads to the formation of orally infective oocysts containing sporozoites (4, 5). For *B. besnoiti*, only tachyzoites and bradyzoites have been described so far. While the genes coding for proteins associated with sexual stages have been identified, sexual reproduction has not been demonstrated and the definitive host, if it exists, is not yet known (3).

*T. gondii* is distributed worldwide and capable of infecting virtually all warm-blooded animals including humans. Transmission occurs through the accidental ingestion of oocysts, or through bradyzoite containing tissue cysts in undercooked meat (6). Vertical transmission by *T. gondii* tachyzoites occurs with maternal primary infection during pregnancy often leading to abortion, stillbirth, or severe developmental abnormalities in the fetus (6, 7), not only in humans, but also in other species such as small ruminants in particular. *T. gondii* infection can also lead to severe neurological and ocular complications, particularly in immunocompromised individuals, and congenitally infected infants. During chronic infection, *T. gondii* bradyzoites encapsulated in tissue cysts are mostly asymptomatic, but the loss of immunocompetence can lead to recrudescence, conversion into tachyzoites, and acute disease. *N. caninum* shares many features with *T. gondii,* among other species primarily affects cattle and dogs, but infections in humans have not been reported (8). Neosporosis in cattle can lead to abortion, birth of weak offspring, or birth of healthy calves that then transmit the parasite to the next generation. In chronically infected animals, pregnancy can lead to recrudescence, and repeated abortions can occur. As a leading cause of reproductive problems in cattle, neosporosis results in substantial economic losses to the livestock industry (4, 9). *B. besnoiti* is less well-studied, and cattle represent the main intermediate host, which suffers from acute disease caused by the rapid proliferation of tachyzoites primarily in endothelia cells of the blood vessels and other cells of mesenchymal origin. The chronic stage is characterized by the presence of large tissue cysts with a multilayered cyst wall, that are formed mostly in the dermis, sclera and mucosa of affected animals. Bovine besnoitiosis is commonly associated with abortion and decreased milk production, weight loss, reduced bull fertility, reduction of hide quality and sporadic abortion due to fever during the acutre phase of infection (10). However, contradictory observations have been reported, and it is clear that there has been insufficient in-depth study of the impact of the disease on production (11, 12). The only mode of transmission known to date is mechanically, either through direct contact from infected animals (e.g. during natural mating), or via flies or biting insects (13). There is no evidence for vertical transmission of *B. besnoiti* to date.

For all three species, the formation of tissue cysts in intermediate hosts is crucial for persistence and disease transmission. *T. gondii* and *N. caninum* typically form cysts in the neural or muscular tissues (5, 14), while *B. besnoiti* forms cysts in the dermis, sclera and mucosal tissues (3). The differentiation into persistent tissue cysts that are largely refractory to drugs is also one of the reasons while current treatment options fail to cure the diseases caused by these parasites. The first-line treatments for acute toxoplasmosis consist of pyrimethamine-sulfadiazine or trimethoprim-sulfamethoxazole that interfere in the folic acid pathway. Other treatment options include the macrolide antibiotic spiramycin (early in pregnancy) and the lincosamide clindamycin (15), or the napthoquinone atovaquone (16). All these drugs are repurposed and lack optimal efficacy and safety profiles. No drugs have been licensed for the use against neosporosis or besnoitiosis so far, although a wide range of compounds have been repurposed in experimental settings (4, 17).

More recently, bumped kinase inhibitors (BKIs), which were either designed on a pyrazolopyrimindine (PP) or 5-aminopyrazole-4-carboxamide (AC) scaffold, have emerged as promising pre-clinical drugs for the treatment of diseases caused by a wide range of apicomplexan parasites (18–22). BKIs were originally designed to selectively target calcium-dependent protein kinase 1 (CDPK1), which is conserved among apicomplexans but absent in mammalian hosts. CDPK1 is required for microneme secretion, gliding motility, host cell invasion and egress (23). More recently it has been shown that BKIs also target *T. gondii* mitogen-activated protein kinase-like 1 (TgMAPKL-1). TgMAPKL1 plays a key role in the *T. gondii* cell cycle by regulating centrosome duplication, which is essential for proper division into two daughter cells during endodyogeny. Mutations in TgMAPKL1 can disrupt this process, leading to defects in cell division and parasite growth (24, 25). Treatments of *T. gondii, N. caninum* and *B. besnoiti* cultured in fibroblasts or MARC cells with a range BKIs resulted not only in impaired host cell invasion, but also in the transformation of intracellular tachyzoites into schizont-like multinucleated complexes (MNCs) (18, 21, 26, 27). These MNCs were characterized by ongoing nuclear division with incomplete cytokinesis and the presence of newly formed intracellular zoites, neither bradyzoites nor tachyzoites, that are not individualized and lack an outer plasma membrane. MNCs remained viable *in vitro* for extended periods of time yet remained confined to the host cell interior. Upon removal of the drug, the effect was reversible, leading to the re-emergence of tachyzoites. MNCs were named “baryzoites”, from the Greek βαρύς = massive, bulky, heavy, or inert (28). Thus, baryzoites constitute a distinct drug-induced stage that promotes parasite survival upon prolonged exposure to elevated drug concentrations (27).

One of the compounds that induced the conversion of tachyzoites into baryzoites is the AC compound BKI-1708 (29). BKI-1708 efficiently inhibited the proliferation of *T. gondii* and *N. caninum in vitro.* In experimentally infected pregnant mice, BKI-1708 treatment was safe and resulted in significantly decreased cerebral parasite loads in the dams, dramatically increased pup survival and strongly reduced vertical transmission (29). In this study, we comparatively analyzed the BKI-1708 induced baryzoites of the closely related cyst-forming coccidians *T. gondii*, *N. caninum*, and *B. besnoiti,* highlighting similarities and species-specific differences on the ultrastructural and proteomic level.

## Materials and methods

### Cell culture media, biochemicals, BKI-1708

Culture medium is from Gibco-BRL (Zürich, Switzerland), and biochemicals are from Sigma (St. Louis, MO, USA). BKI-1708 was originally synthesized in the Department of Biochemistry of the University of Washington, USA and scaled up by WuXi Apptec Inc., Wuhan, China to >98% purity by LC/MS-MS and NMR and shipped as powder (30). For *in vitro* studies, stock solutions of 20 mM were prepared in dimethyl-sulfoxide (DMSO) and stored at −20 °C.

### Host cells and parasites

Human foreskin fibroblasts (HFF; PCS-201-010™) were maintained as previously described (28). The strains used in this study were *T. gondii* ME49, *N. caninum* Spain-7 isolate (NcSpain-7) and *B. besnoiti* Bblis14. They were maintained in HFF as previously described in (21, 31, 32).

### BKI-1708 treatments and transmission electron microscopy (TEM)

For TEM, HFF were grown to confluence in T25 flasks in culture medium for three days at 37 °C/5 % CO_2_ and were infected with 1 × 10⁶ *T. gondii* ME49, *N. caninum* Nc-Sp7 or *B. besnoiti* Bblis14 tachyzoites. At 4h post-infection, the medium was supplemented with 2.5 µM BKI-1708. The medium was removed after 24 hours for untreated cultures, and at days 4 and 6 of continuous treatment, with BKI-1708 supplemented medium being renewed every second day. At the desired timepoints, cultures were washed once with 100 mM sodium cacodylate, pH 7.3, and fixation was carried out in 100 mM cacodylate buffer containing 2 % glutaraldehyde for 10 min. Then, adherent cells were carefully removed with a cell scraper and transferred into a tube. After another 2-4 hours of fixation at room temperature, samples were centrifuged and post-fixed in 2 % cacodylate buffer containing 2 % osmium tetroxide for 2 hours. Following several washes in distilled water, specimens were dehydrated through a graded series of ethanol (30, 50, 70, 90 and 3 x 100%), and resuspended in Epon 812 epoxy resin as previously described (33). After 3 changes of Epon resin, polymerization was carried out at 60 °C overnight. Sections of 80 nm thickness were cut on an ultramicrotome (Reichert and Jung, Vienna, Austria) and were transferred onto formvar-carbon-coated 200 mesh nickel grids (Plano GmbH, Marburg, Germany). They were stained with Uranyless® and lead citrate (Electron Microscopy Sciences, Hatfield PA, USA), and specimens were inspected on a FEI Morgagni TEM equipped with a Morada digital camera system (12 Megapixel) operating at 80 kV.

### Reversion and Long-Term Treatment Assays

5 × 10⁵ HFF were grown to confluence in T25 flasks. These monolayers were then infected with 5 × 10⁶ tachyzoites of TgME49, Nc-Sp7, or Bblis14 tachyzoites. Cultures were washed with phosphate buffered saline (PBS) at 1h post-infection and were exposed to 2.5 µM BKI-1708 during 5 days at 37°C / 5% CO_2_. BKI-supplemented media was renewed on the third day of treatment, and control cultures were left untreated. After 5 days of continuous treatment, cultures were washed with PBS and further maintained in fresh medium without drug. Regrowth of parasites cultures was monitored every day by conventional light microscopy until host cell lysis (lysis plaque formation) and extracellular tachyzoites were visible. To assess the effects of a more prolonged treatment on *T. gondii*, 5 × 10⁵ HFF were grown to confluence in T25 flasks and subsequently infected with 5 × 10⁶ *T. gondii* ME49 tachyzoites. One hour post infection, cultures were washed with PBS and the medium was supplemented with 2.5 µM BKI-1708, followed by culture at 37 C / 5 % CO_2_for 30 days. Cultures were left in BKI-supplemented medium and monitored every second day with conventional light microscopy.

### Comparative proteomics

Semi-confluent HFF monolayers were maintained in T75 cell culture flasks, and were infected with 1 x 10⁷ TgME49, Nc-Spain7 or Bblis14 tachyzoites. At 4 h post infection, treatment with 2.5 µM BKI-1708 was initiated, control cultures were not exposed to BKI-1708. Three biological replicates were used for each. All non-treated control cultures were maintained at 37 ◦C/5% CO_2_ during 3 days, BKI-1708 treated cultures during 5 days (BKI-1708). Subsequently, the infected monolayers were washed twice with PBS, followed by removal of infected cells with a rubber cell scraper and resuspension in PBS followed by centrifugation (15 min, 1000× g, 4 ◦C). Cell pellets were lysed in 100 µL 8M urea/100 mM Tris/HCl pH 8/cOmpleteTM protease inhibitor cocktail (Roche Diagnostics, Rotkreuz, Switzerland) by incubation for 15 min at RT followed by 15 min in an ultrasonic water bath. Proteins were reduced and alkylated with 10 mM DTT for 30 min at 37 ◦C and 50 mM iodoacetamide for 30 min at 37 ◦C. Proteins were precipitated at −20 ◦C by addition of 5 volumes cold acetone and incubation at −20 ◦C overnight. All liquid was carefully removed, and the pellet dried in ambient air for 15 min before reconstitution of proteins in 8 M urea, 50 mM Tris-HCl pH 8.0 to an approximate protein concentration of 2 mg/mL. Effective protein concentrations were determined by Bradford assay (1:10 diluted samples) and 10μg of proteins were digested and analyzed by shotgun nano liquid chromatography coupled to tandem mass spectrometry as described elsewhere (34).

The mass spectrometry data was searched and quantified with Spectronaut (Biognosys) version 19.0.240606.62635 in the hybrid directDIA+ (Deep) with factory settings. Search parameters included Acetyl (Protein N-term) and Oxidation (M) as variable modifications (5 max variable modifications), Carbamidomethyl (C) as fixed modification, Trypsin/P (max 2 missed cleavages) as digestion enzyme. The Tg+host, Nc+host and Bb+ host samples were searched respectively against the ToxoDB-55 (35). *T. gondii* ME49 annotated protein sequences, a concatenation of swissprot and uniport *N. caninum* (strain Liverpool) sequences (https://www.uniprot.org; release July 2024), and a concatenation of swissprot and ToxoDB-68 *B. besnoiti* sequences. Common contaminants were added in all 3 databaes. Protein groups with less than 2 peptides per group were excluded. A leading protein was chosen per protein group on the basis of best coverage, and the IBAQ (Intensity-based absolute quantification) values calculated by Spectronaut are reported here with respect to the leading protein. A relative abundance (rAbu) was calculated so that the sum of rAbu were 1000000 for every sample.

Protein groups not flagged as potential contaminants and for which we had at least 2 detections in at least one of the replicate groups were considered for differential expression. Missing values were then imputed in the following manner: if there was at most two non-zero values in the replicate group then the missing values were imputed by a left-censored method; this was done by drawing random values from a Gaussian distribution of width 0.3 x sample standard deviation centered at the sample distribution mean minus 2.5 x sample standard deviation. Any remaining missing values were imputed by the Maximum Likelihood Estimation (MLE) method (36) Differential expression tests were performed with the moderated t-test of the R limma package (37). The Benjamini and Hochberg (38) method was then applied to correct for multiple testing. The criterion for statistically significant differential expression was that the largest accepted adjusted p-value reaches 0.05 asymptotically for large absolute values of the log2 fold change and tends to 0 as the absolute value of the log2 fold change approaches 1 (with a curve parameter of 0.1x overall standard deviation). Proteins consistently significantly differentially expressed through 20 imputation cycles were flagged accordingly (39).

### Fluorescence microscopy

For immunofluorescence studies, 2 × 10^4^ HFF were grown on glass coverslips in 24-well plates at 37 °C and 5% CO_2_ for three days, and then infected with 2 × 10^4^ TgMe49, NcSpain-7 or Bblis14 tachyzoites. Four hours after infection, the medium was replaced and supplemented with 2.5 μM BKI-1708, or medium without compound. Cultures were maintained at 37 °C and 5% CO_2_ for a maximum of 9 days, with BKI-1708 supplemented medium being changed every second day. On days 2, 6 and 9, cells were fixed in 3% paraformaldehyde in PBS (pH 7.2) for 10 min, and permeabilized in a 1:1 solution of pre-cooled methanol/acetone at −20 °C. Following rehydration, nonspecific binding sites were blocked in PBS/3% bovine serum albumin (BSA) overnight at 4 °C.

For immunofluorescence double staining, primary and secondary antibodies were diluted in PBS containing 0.3% BSA and added sequentially to the samples during 30 minutes each. The primary antibodies used in this study are listed in Table 1, corresponding secondary antibodies were tetramethyl-rhodamine-isothiocyanate (TRITC)-conjugated anti-mouse, FITC-conjugated anti-rabbit, and TRITC-conjugated anti-bovine serum all at a dilution of 1:300. Following antibody staining, samples were thoroughly washed in PBS and were mounted in, H-1200 Vectashield mounting medium with DAPI (4′,6-diamidino-2-phenylindole; Vector Laboratories, Inc. Burlingame, CA, USA), and viewed on a Nikon E80i fluorescence microscope. Processing of images was performed using ImageJ software from National Institutes of Health, Bethesda, MD, USA (40).

**Table 1.**
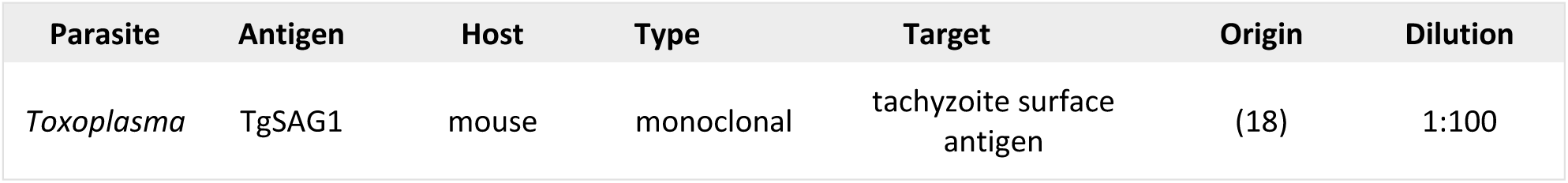
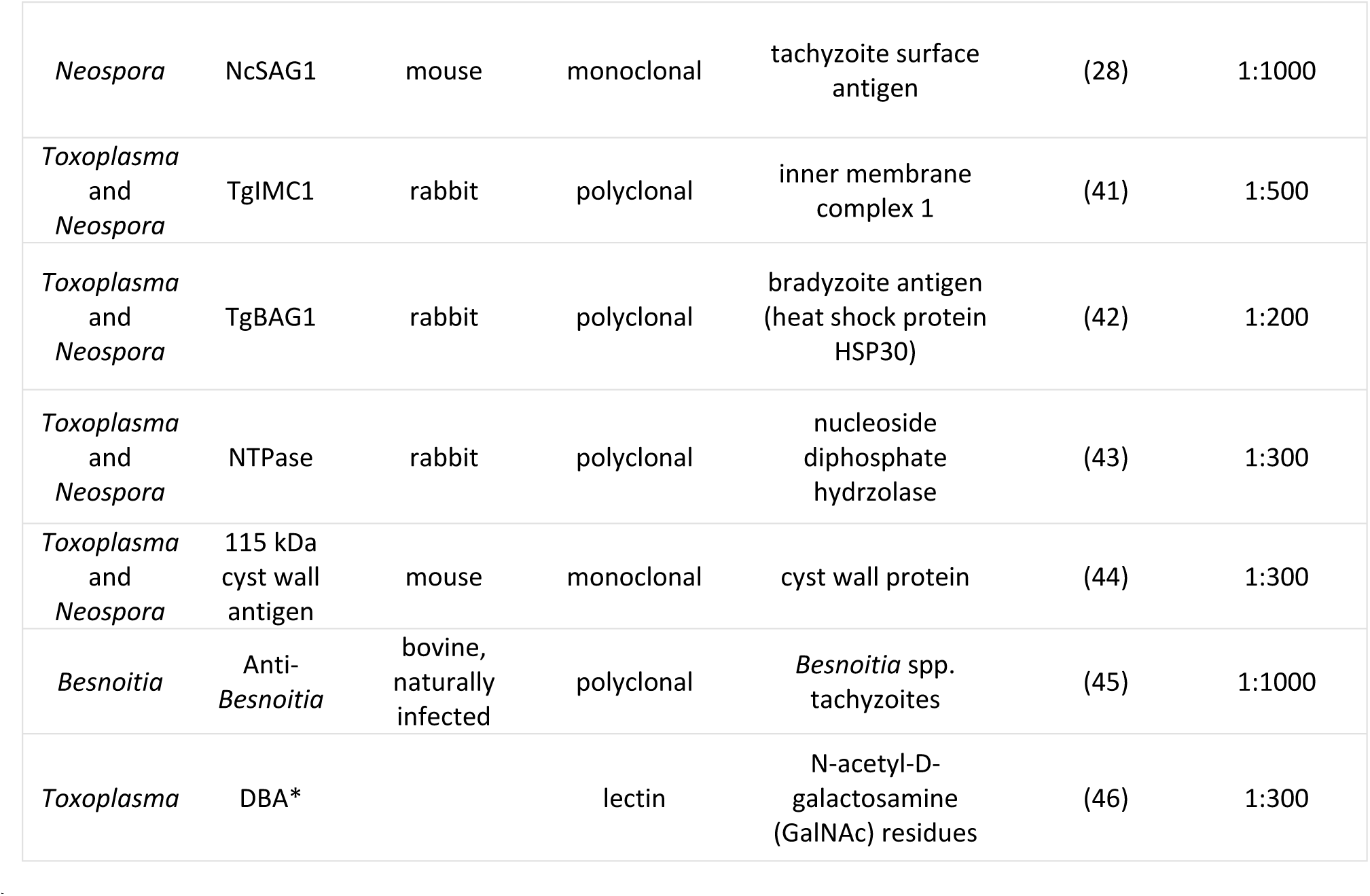
Primary antibodies used for immunofluorescence (IF). Host species, antibody type, target, origin and working dilutions are indicated. \**Dolichos Biflorus* Agglutinin (DBA) is a lectin conjugated to fluorescein isothiocyanate (FITC).

| Parasite | Antigen | Host | Type | Target | Origin | Dilution |
| --- | --- | --- | --- | --- | --- | --- |
| *Toxoplasma* | TgSAG1 | mouse | monoclonal | tachyzoite surface antigen | (18) | 1:100 |
| *Neospora* | NcSAG1 | mouse | monoclonal | tachyzoite surface antigen | (28) | 1:1000 |
| *Toxoplasma*<br>and<br>*Neospora* | TgIMC1 | rabbit | polyclonal | inner membrane complex 1 | (41) | 1:500 |
| *Toxoplasma*<br>and<br>*Neospora* | TgBAG1 | rabbit | polyclonal | bradyzoite antigen (heat shock protein HSP30) | (42) | 1:200 |
| *Toxoplasma*<br>and<br>*Neospora* | NTPase | rabbit | polyclonal | nucleoside diphosphate hydrolase | (43) | 1:300 |
| *Toxoplasma*<br>and<br>*Neospora* | 115 kDa cyst wall antigen | mouse | monoclonal | cyst wall protein | (44) | 1:300 |
| *Besnoitia* | Anti-*Besnoitia* | bovine, naturally infected | polyclonal | *Besnoitia* spp. tachyzoites | (45) | 1:1000 |
| *Toxoplasma* | DBA* |  | lectin | N-acetyl-D-galactosamine (GalNAc) residues | (46) | 1:300 |

## RESULTS

### Ultrastructural features of *T. gondii, N. caninum* and *B. besnoiti* baryzoites induced by treatment with BKI-1708

Figure 1 displays electron micrographs of *T. gondii, B. besnoiti* and *N. caninum* tachyzoites grown in HFF monolayers in the absence of BKIs. The three species exhibited a largely similar ultrastructure. Tachyzoites reside and proliferate within a parasitophorous vacuole that is delineated by a parasitophorous vacuole membrane (indicated by red arrows in Fig 1). They displayed the typical apicomplexan features including an apical complex with the conoid, and secretory organelles such as rhoptries, micronemes and dense granules. Depending on the section plane, larger or smaller portions of the single mitochondrion were visible, including the mitochondrial matrix with cristae of varying electron density. Tachyzoites divided by endodyogeny with two daughter zoites being formed that represented clearly separate entities. Frequently, daughter tachyzoites were seen still attached to a residual body.

**Figure 1.**
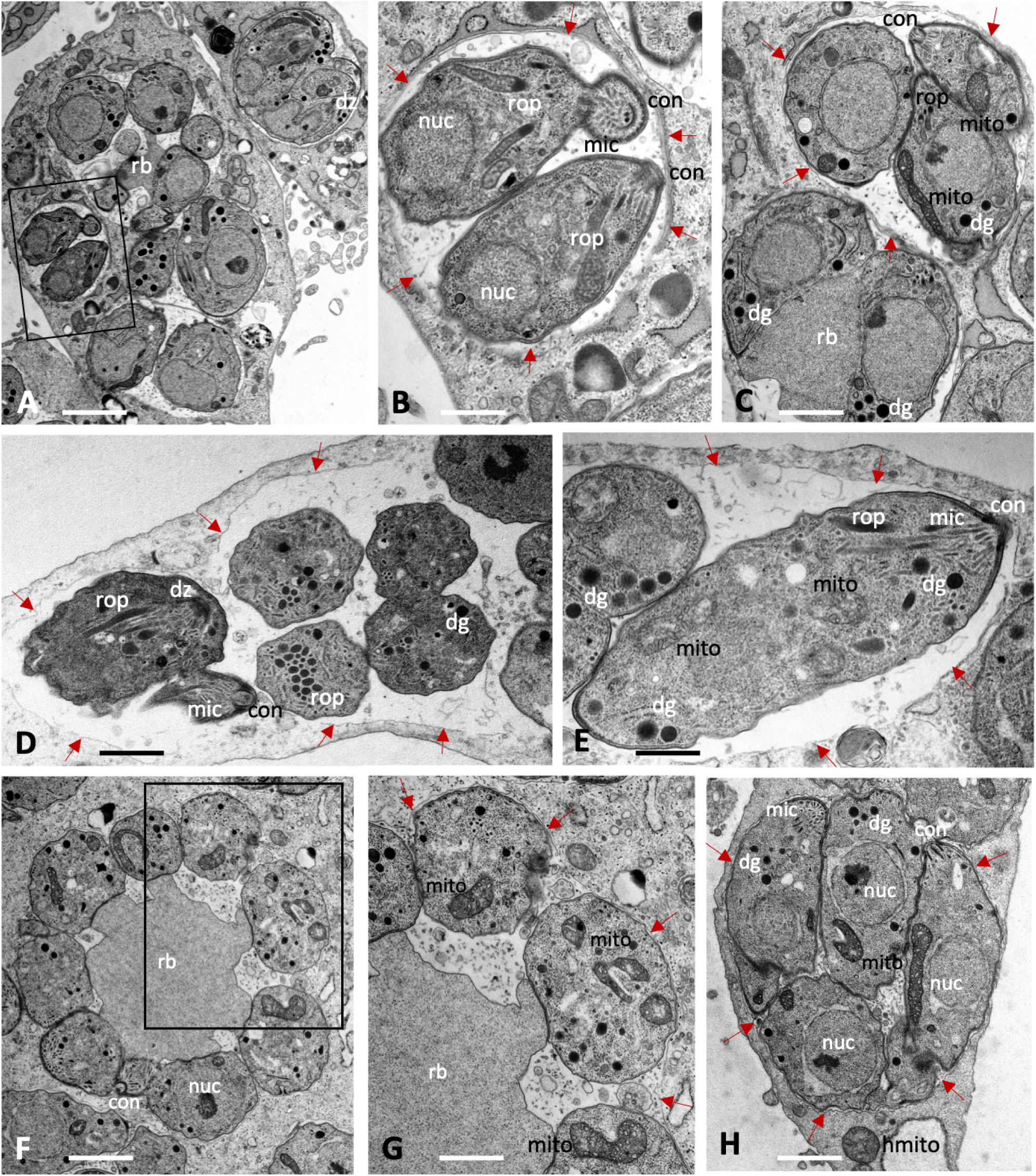
Tachyzoites of *Toxoplasma gondii* (A-C), *Neospora caninum* (D, E) and *Besnoitia besnoiti* (F-H) maintained in human foreskin fibroblasts for 48 h in the absence of compound treatment. The boxed areas in A and F are enlarged in B and G, respectively. Tachyzoites undergo intracellular proliferation in a parasitophorous vacuole, which is delineated by a parasitophorous vacuole membrane (marked with red arrows); nuc = nucleus, rop = rhoptries, mic = micronemes, dg = dense granules, mito = mitochondrion, con = conoid, rb = residual body, dz = newly formed daughter zoites. Bars in A = 2.3 µm; B = 0.75 µm; C = 0.85 µm; D = 1.1 µm; E = 0.75 µm, F = 1.25 µm; G = 0.75 µm; H = 1.1 µm

Treatment with BKI-1708 of *Toxoplasma gondii* ME49 cultured in HFF led to the transformation of intracellular tachyzoites into schizont-like and thus multinucleated baryzoites, while single tachyzoites were not detected (Figure 2). *T. gondii* baryzoites formed after 4 days of treatment are shown in Fig 2A-C, and baryzoites formed after 6 days of BKI-1708 treatment are shown in Figs 2D-G. Baryzoites contained numerous nuclei that form clusters either in the periphery or to one side of the cytoplasm, depending on the section plane. Small apical complexes of daughter zoites were frequently seen protruding at the periphery or within the cytoplasmic areas, which also contain the typical organelles reminiscent for apicomplexans, including dense granules, rhoptries, as well as portions of a structurally intact mitochondrion including an electron-dense matrix and cristae. However, these newly formed zoites lacked the typical triple layered membrane of tachyzoites. Instead, this membrane covered the entire baryzoite surface. *T. gondii* baryzoites were seen to release large amounts of secretory vesicles and lamellar membranous components into the matrix of the parasitophorous vacuole. At 6 days of treatment it appeared that at least some of this material formed electron dense deposits at the periphery of the vacuole (Figs 2F and G). During this remarkable drug-induced transformation, however, based on ultrastructural appearance, parasites were largely viable.

**Figure 2.**
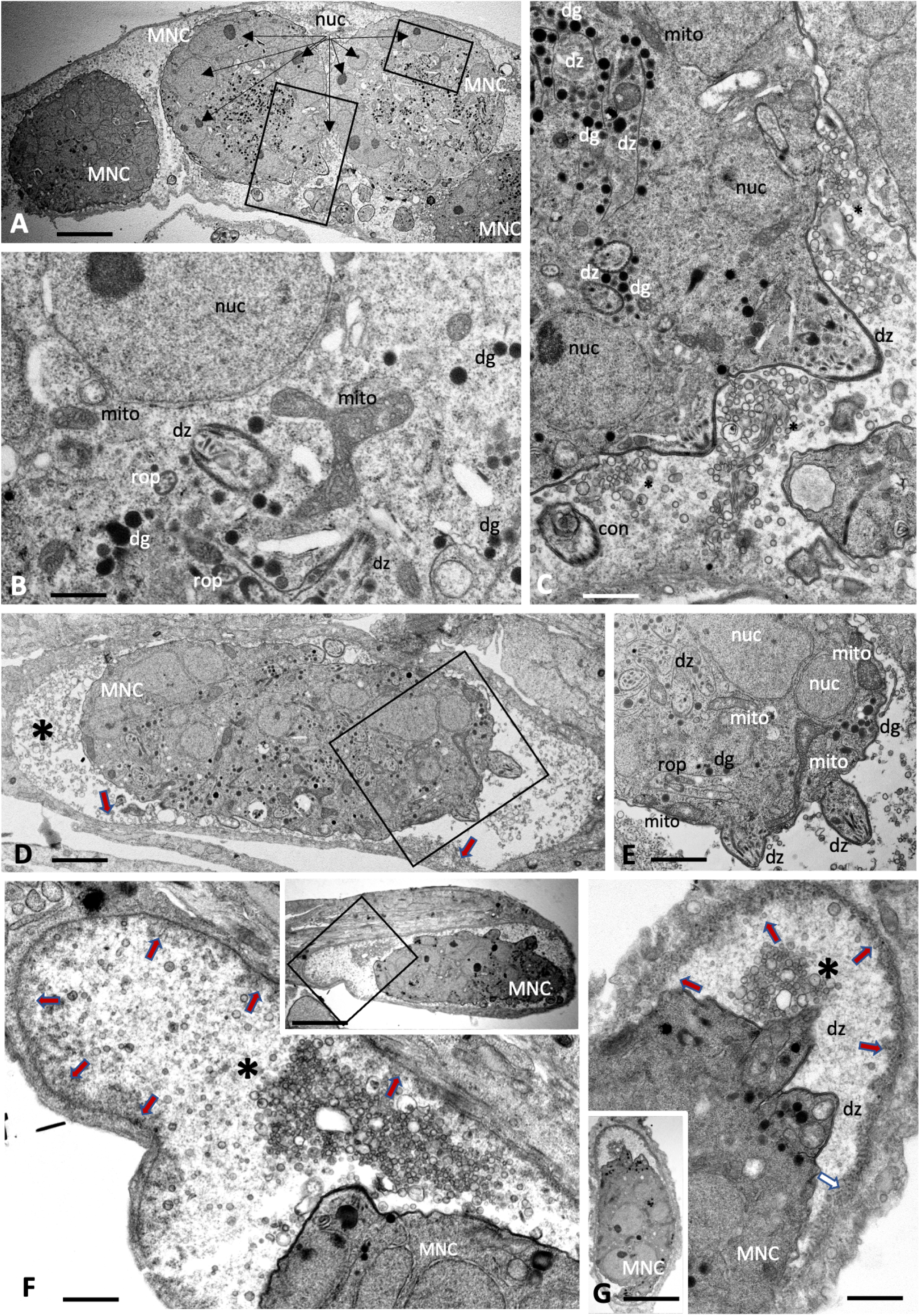
Baryzoites of *T. gondii* grown in human foreskin fibroblast cultures treated with BKI-1708 during 4 (A-C) and 6 days (D-G). The boxed areas in A are shown at higher magnification in B and C, the one in D is magnified in E. Lower magnifications views of F and G are shown in the corresponding inserts. MNC = multinucleated complex, nuc = nucleus, rop = rhoptries, dg = dense granules, mito = mitochondrion, dz = apical parts of daughter zoites, * indicates secretory products in the parasitophorous vacuole. The red arrows in D, F and G point towards the developing cyst wall. Bars in A = 3.2 µm; B = 0.5 µm; C = 0.75 µm; D = 3.2 µm; E = 1.7 µm; F = 1.2 µm; F insert = 6.7 µm; G = 1.5 µm; G insert = 6.7 µm.

Baryzoites formed upon exposure of *N. caninum* to BKI-1708 (Figure 3) exhibited largely similar characteristics and also remained within their host cell. The matrix of the parasitophorous vacuole was composed of rather evenly distributed vesicular and granular components, but an electron-dense cyst wall like structure was clearly missing. As in *T. gondii,* nuclei were clustered together and separated from other cytoplasmic compartments of newly formed daughter zoites (Figs 3C, F, G, H), and mitochondria also remained structurally intact (Figs 3D, E, G, H), indicating that there is no impairment of viability. Compared to *T. gondii* baryzoites, however, *N. caninum* baryzoites contained an increased number of organelles that resembled dense granules (dg) exhibiting variable diameters (150-400 nm). Occasionally lipid droplets (ld) were seen (Fig. 3 A, G). In addition, an increased number of vacuolar inclusions of varying sizes were observed that either appeared empty or contain undefined granular or filamentous material or electron dense deposits, the nature of which is unknown (Fig 3C-H).

**Figure 3.**
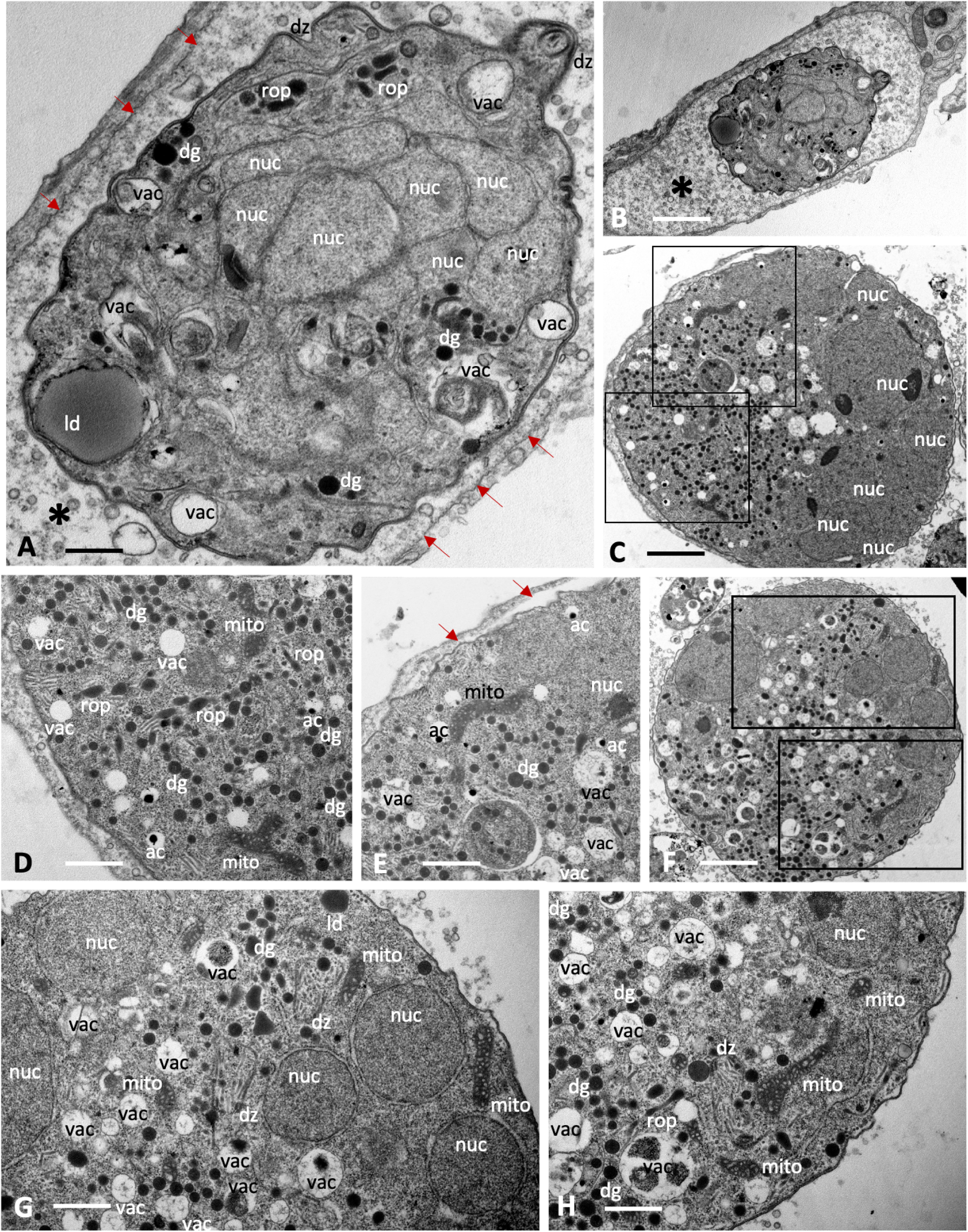
Baryzoites of *N. caninum* grown in human foreskin fibroblast cultures treated with BKI-1708 during 4 (A-C) and 6 days (D-H). The MNC in A is shown enlarged in B, the * marks the interior matrix of the parasitophorous vacuole, red arrows point towards the parasitophorous vacuole membrane. Higher magnification views of the boxed areas in C are depicted in D and E. Selected areas in F are enlarged in G and H; nuc = nucleus, dg = dense granules, rop = rhoptries, vac = cytoplasmic vacuoles, ld = lipid droplets, mito = mitochondrion, ac = acidocalcisome, dz = apical part of a daughter zoite. Bars in A = 2.3 µm; B = 0.65 µm C = 1.6 µm; D = 0.8 µm; E = 0.9 µm; F = 1.6 µm; G = 0.8 µm; H = 0.8 µm

*B. besnoiti* baryzoites exhibited similar features, with nuclei and cytoplasmic components largely compartmentalized (Figure 4). However, the degree of vacuolization appeared higher compared to *T. gondii* and *N. caninum*, including also irregularly shaped vacuoles. In many baryzoites, the central portion of the mitochondrial matrix appeared slightly distorted, but more electron dense in the periphery (Fig 4F). Generally, the typical apicomplexan secretory organelles such as micronemes, rhoptries and dense granules were less well preserved. Numerous dense granules were found in the cytoplasm, albeit of small size (200 nm or below). In *B. besnoiti,* the matrix of the parasitophorous vacuole (indicated with an asterisk) was filled with small secreted vesicles (Figs 4E, F), as well as also larger vesiculated structures (Figs 4G, H), and additionally clusters of electron dense bodies (Figs 4I, J) of unknown composition.

**Figure 4.**
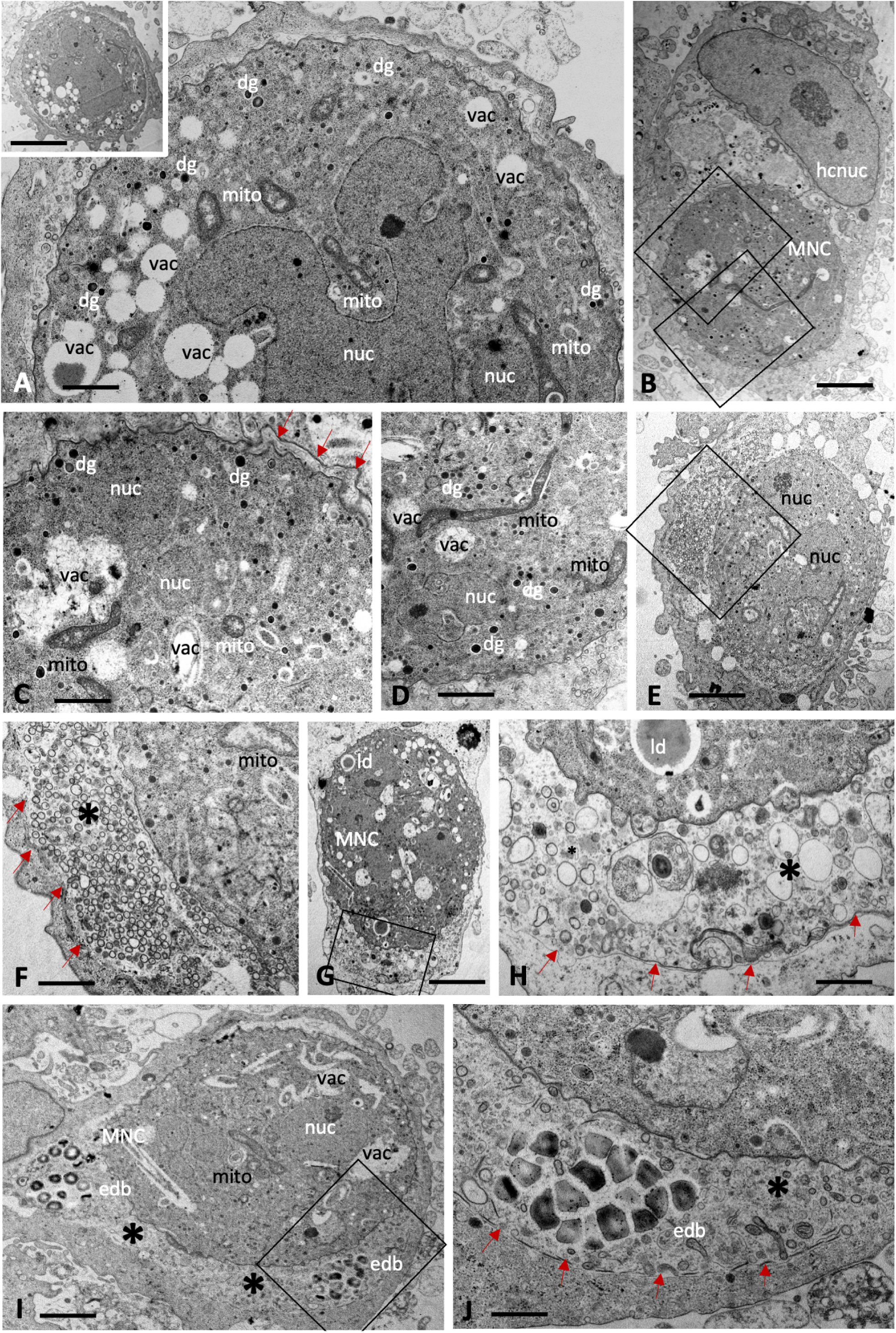
Baryzoites of *B. Besnoiti* grown in human foreskin fibroblast cultures treated with BKI-1708 during 4 (A-E) and 6 days (F-J). The insert in A shows a lower magnification view of the respective MNC, and the two boxed areas in B are enlarged in C and D. F shows an enlarged view of the boxed area in E, and the framed areas in G and I are shown at higher magnification view in H and I. Red arrows point towards the parasitophorous vacuole membrane. MNC = multinucleated complexes, nuc = nucleus, dg = dense granules, hcnuc = host cell nucleus, vac = cytoplasmic vacuoles, edb = electron dense bodies, mito = mitochondrion, ld = lipid droplet. * = matrix of the parasitophorous vacuole, arrows point at parasitophorous vacuole membrane. Bars in A = 0.8 µm; A insert = 4.2 µm; B = 2 µm; C = 0.7 µm; D = 1.2 µm; E = 2 µm; F = 0.7 µm, G = 3.6 µm, H = 0.7 µm, I = 2.4 µm; J = 0.8 µm

### BKI-1708 does not act parasiticidal and removal of BKI-1708 leads to reversion into tachyzoites

Infected cultures were treated with 2.5 µM BKI-1708 for 5 days, followed by culture in the absence of drug treatment and daily microscopical inspection. All the three species were able to recover from treatment and reverted from baryzoites into the infective tachyzoite stage, but they did so at different rates (Fig 5A). *B. besnoiti* baryzoites resumed proliferation more rapidly, with baryzoite-to-tachyzoite reversion observed after 5 days of drug removal. *T. gondii* and *N. caninum* baryzoites required 10 and 9 days, respectively, to revert into infective tachyzoites. In addition, prolonged treatment of *T. gondii* infected cultures for a period of 30 days did not clear the infection or lead to disintegration of the complexes (Fig 5B). Baryzoites remained intact and morphologically similar to those observed after shorter treatment periods.

**Figure 5.**
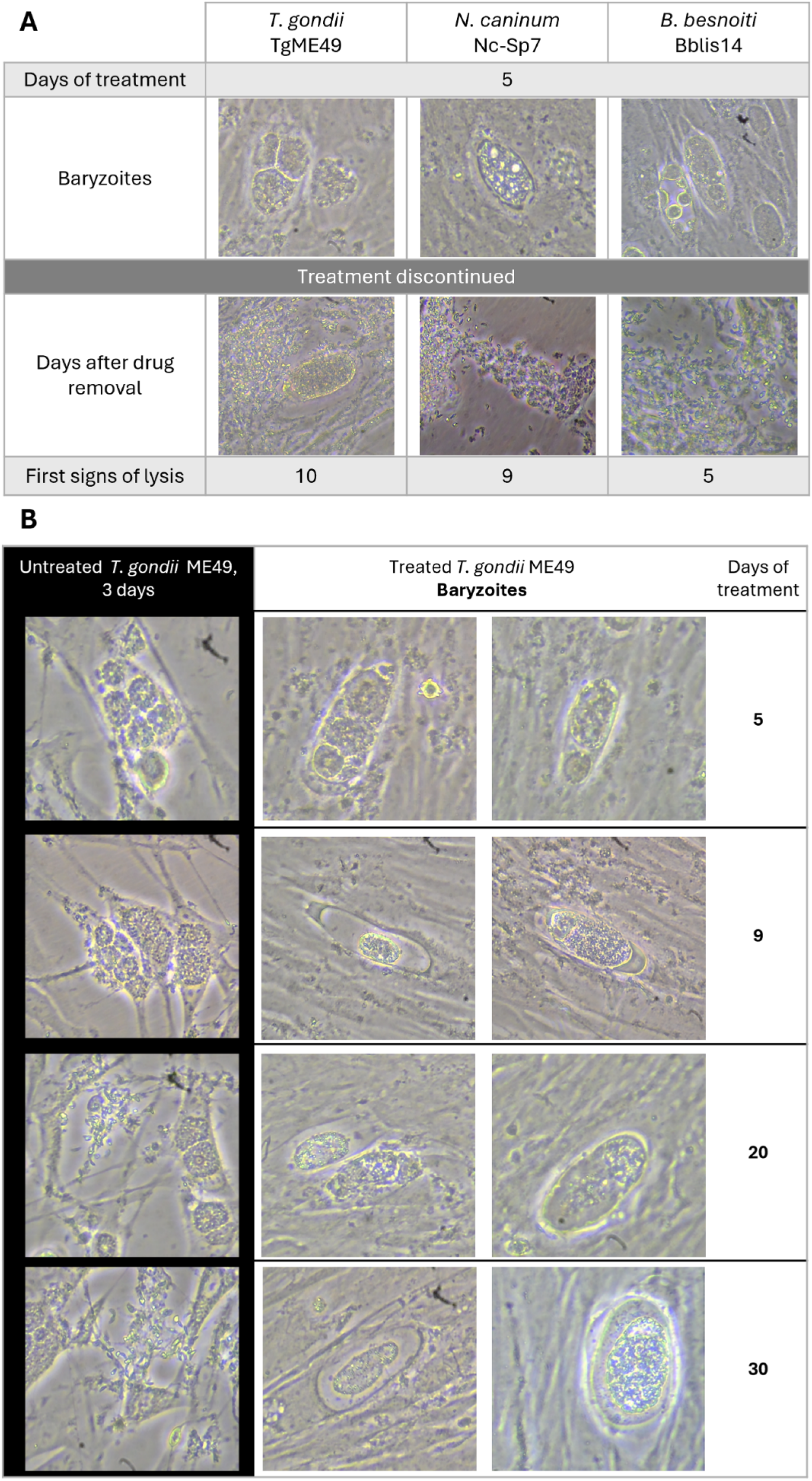
Reversion and Long treatment Assays. (A) Tachyzoites were treated with BKI-1708 at 2.5 μM 1h post-infection for 5 consecutive days, leading to the formation of baryzoites. On day 5, drug pressure was released, and cultures are monitored daily by conventional light microscopy until lysis plaques are detected. Regrowth of parasites was noted in all cases. (B) ME49 tachyzoites were treated for a total period of 30 days and monitored by conventional light microscopy. Intact baryzoites were observed even after 30 days of continuous treatment.

### Upon treatment with BKI-1708, *T. gondii* baryzoites upregulate bradyzoite-specific markers and downregulate tachyzoite antigens

As shown by immunofluorescence in Figure 6, the typical untreated apicomplexan tachyzoites of *T. gondii* and *N. caninum* were labelled with antibodies directed against TgSAG1 and NcSAG1, respectively, and antibodies directed against IMC1. In *T. gondii* treated with BKI-1708, TgSAG1-staining no longer defined the contours of individual tachyzoites, but was found on the surface the developing baryzoites, with diminished staining intensity after 6 days of treatment (Fig 6A). A similar phenotype was observed in *N. caninum* baryzoites, where BKI-1708 treatment also resulted in altered NcSAG1 localization, accumulation of nuclei and IMC1-stained zoites (Fig 6B). Overall, these results mirrored earlier reports on the effects of BKI-1294 in *N. caninum* (28).

**Figure 6.**
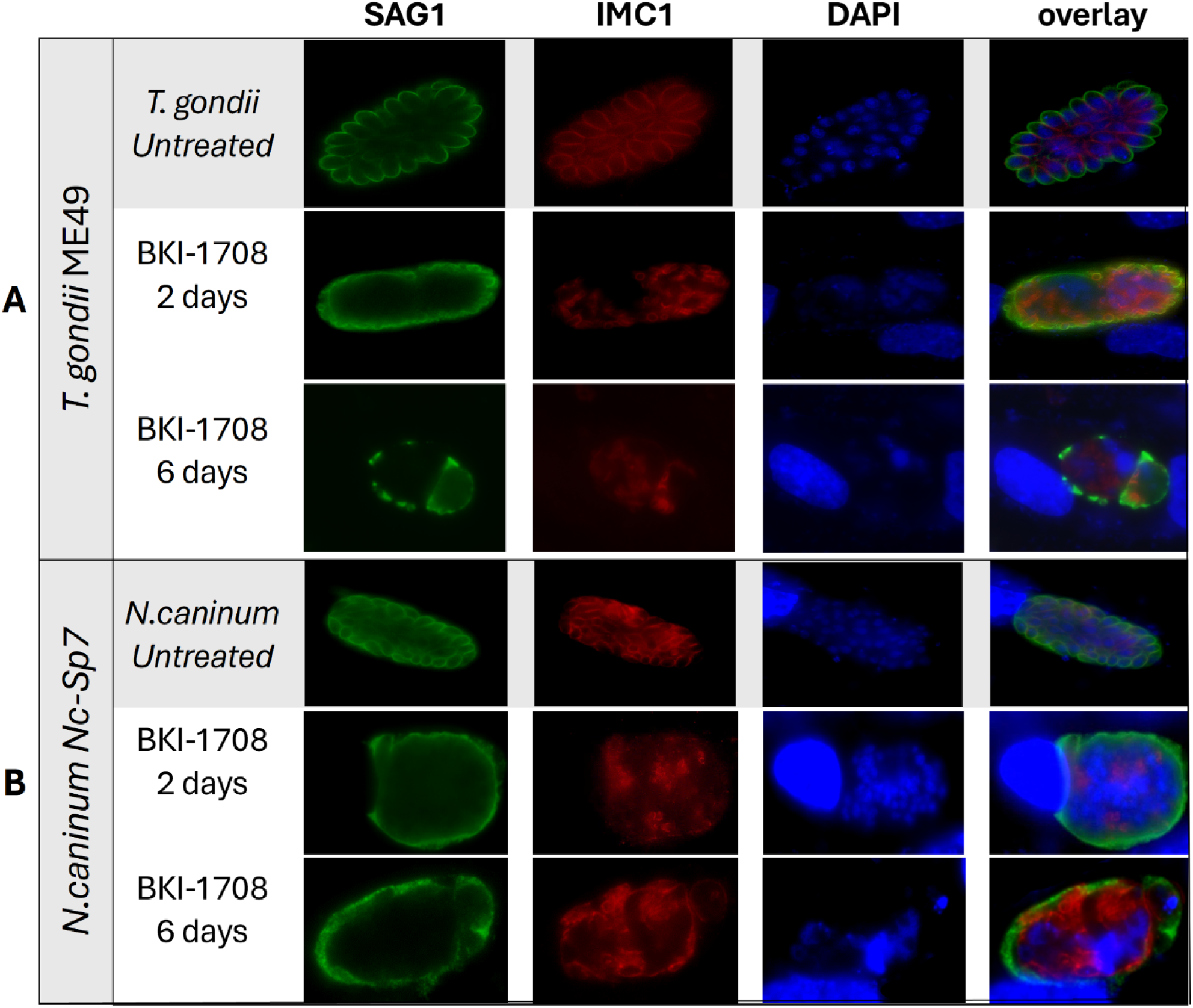
Immunofluorescence staining of *T. gondii* (A) and *N. caninum* (B) untreated (control) and treated tachyzoites (baryzoites) after 2 and 6 days. Green labels the major tachyzoite surface antigen (SAG1); Red delimitates individual zoites, labeling the inner membrane complex I(IMC1); DNA of nuclei is blue, stained by DAPI.

Prolonged exposure to BKI-1708 for 9 days resulted in the disappearance of detectable TgSAG1 staining on the *T. gondii* baryzoites (Figs 7A and C). From 6 days of treatment onwards, *T. gondii* baryzoites expressed the bradyzoite marker TgBAG1) (Fig 7A). In addition, *T. gondii* baryzoites switched on the expression of cyst wall associated markers recognized by mAb CC2 and DBA (Figs 7B and C). However, in *N. caninum* baryzoites, NcSAG1 expression on the baryzoite surface remained unaltered even after 9 days of BKI-1708 exposure, while no labeling was detected using the cyst wall markers mAbCC2 and DBA (Figs 7D and E). Antibodies directed against TgBAG1 did also not react with *N. caninum* baryzoites (data not shown) and none of the fluorescent markers stained *B. besnoiti* baryzoites (Fig 7F).

**Figure 7.**
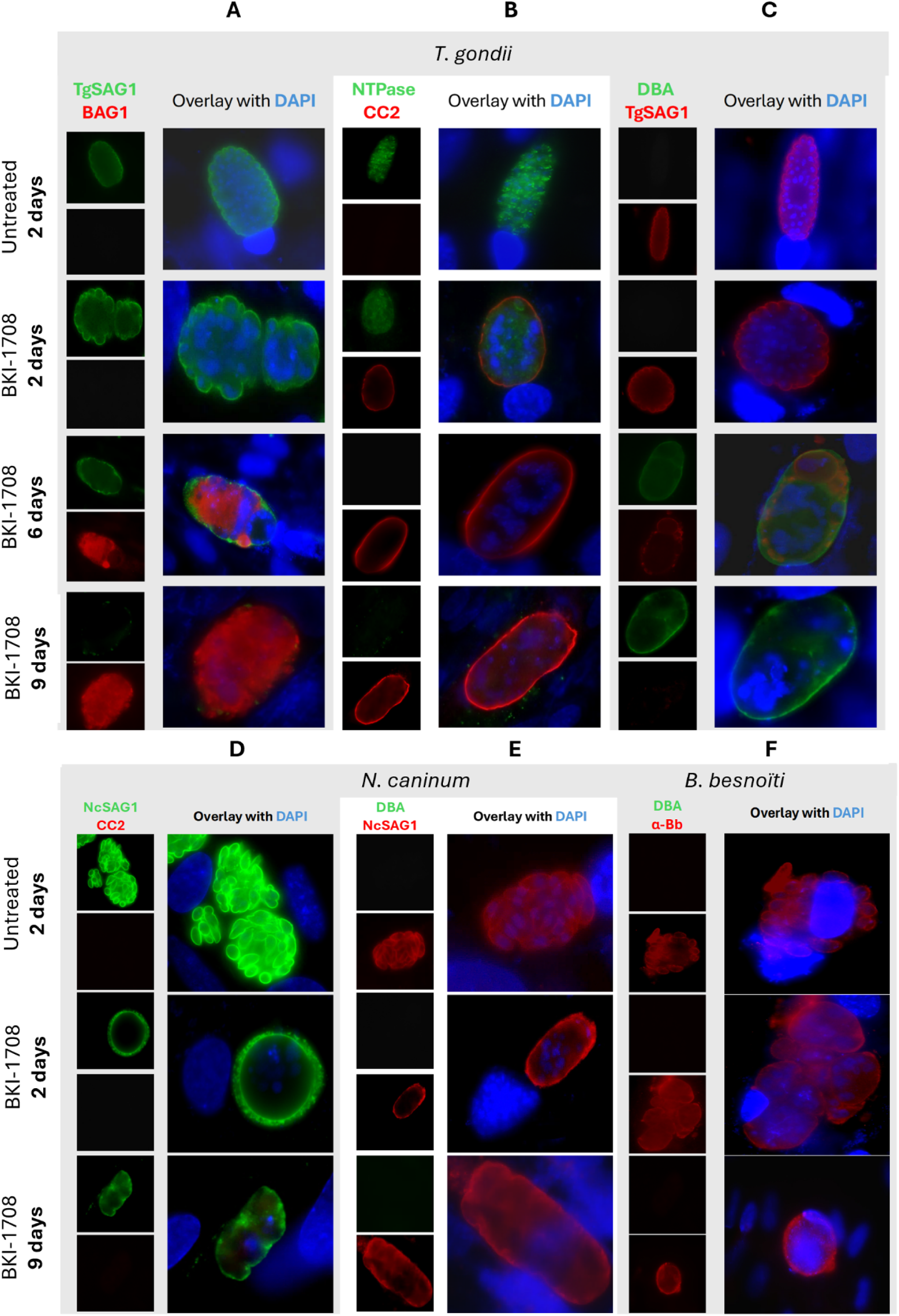
Immunofluorescence of *T. gondii* (A-C), *N. caninum* (D, E) and *B. besnoiti* (C) under BKI-1708 treatment. (A) Major surface antigen SAG1 is stained in green; Red labels bradyzoite cytoplasmic proteins (BAG1); (B) Nucleoside-triphosphatase (NTPase) enzyme is labeled green; α-CC2 staining is red; (C) Dolichos biflorus agglutinin (DBA) is stained with green; Major surface antigen SAG1 is stained in red; (D) NcSAG1 major antigen stained in green, CC2 labelled in red; E) NcSAG1 stained in red, DBA is labeled in green; Anti-Besnoitia spp. staining is red, and DBA is labeled green. Nuclei in all panels were stained with DAPI (blue).

### Comparative proteomics study of BKI-1708 treated baryzoite cultures reveals differentially expressed proteins in *T. gondii, N. caninum and B. besnoiti*

*T. gondii*, *N. caninum* and *B. besnoiti* infected HFF were treated with BKI-1708 for 5 days and were subjected to comparative proteomic analyses, thus revealing the presence of differentially expressed proteins in baryzoites compared to untreated tachyzoites. Overall, 2670 proteins were identified in cultures infected with *T. gondii* baryzoites, 1461 in *N. caninum*, and 2755 proteins were identified in *B. besnoiti* baryzoite cultures (Table 2). Unbiased analysis of the dataset by principal component analysis (PCA) of the log2-transformed spectronaut (SN) intensities of the samples demonstrated that the proteomes of tachyzoites and BKI-1708 induced baryzoites formed non-overlapping clusters separated by both principal components (Figure 8A). In addition, SN protein intensity distributions were almost equal for all samples (Figure 8B). The entire proteomic datasets for *T. gondii*, *N. caninum* and *B. besnoiti* are presented in Supplementary Tables S1, S2 and S3, respectively.

**Figure 8.**
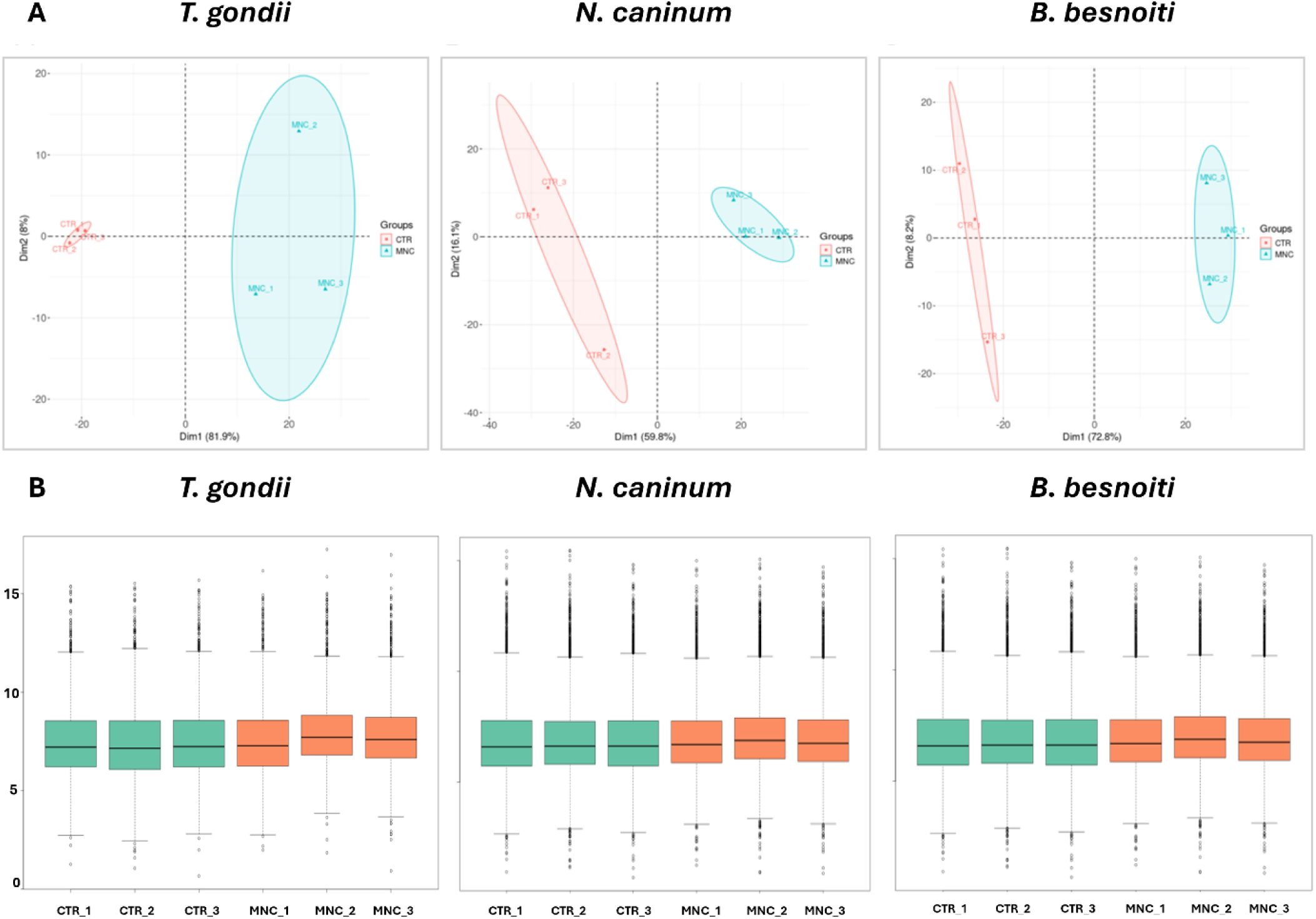
Proteomic profiling of multinucleated complexes (baryzoites) in *Toxoplasma gondii*, *Neospora caninum*, and *Besnoitia besnoiti*. (A) Principal component analysis (PCA) of log₂-transformed SN intensities from proteome datasets of untreated tachyzoites (CTRL) and treated parasites forming multinucleated complexes (MNC) for *T. gondii* ME49, *N. caninum* Nc-Sp7, and *B. besnoiti* Bblis14. (B) Box plots of log₂-transformed leading SN intensities of the same proteome datasets comparing untreated tachyzoites (CTRL) and MNC for the three species. MNC refers to multinucleated complexes or baryzoites.

**Table 2.** Summary of protein abundance changes in *T. gondii* ME49, *N. caninum* Nc-Sp7, and *B. besnoiti* BbLis14 baryzoites compared to untreated tachyzoites.

|  | *T. gondii*<br>baryzoites | *N. caninum*<br>baryzoites | *B. besnoiti*<br>baryzoites | Commonly<br>DE proteins |
| --- | --- | --- | --- | --- |
| Total number of detected proteins | 2670 | 1461 | 2755 | 19 |
| Number of detected proteins in<br>higher abundance | 153 (5,7%) | 15 (1%) | 39 (1,4%) | 3 |
| Number of detected proteins in<br>lower abundance | 290 (10,9%) | 114 (7,8%) | 389 (14,1%) | 16 |

The basic hallmarks of the three proteomes and respective differentially expressed (DE) proteins are depicted in Table 2. For all three species, the number of baryzoite proteins with downregulated abundance compared to tachyzoites was higher than the number of proteins with upregulated expression levels. *B. besnoiti* baryzoite cultures exhibited the strongest downregulatory response to treatment, with 14.1 % of all proteins expressed at lower abundance, compared to 10.9 % of *T. gondii* and 7.8 % of *N. caninum* proteins. Conversely, upon BKI-1708 treatment, the largest proportion of proteins was upregulated in *T. gondii* baryzoites (5.7%) compared to *B. besnoiti* and *N. caninum* baryzoites (1.4 and 1%, respectively). A total of sixteen orthologs were commonly found in all three species at lower abundance in baryzoite cultures, while three were consistently found to be expressed at higher levels compared to untreated tachyzoites in all three species (Table 2).

The DE proteins of all three species were grouped into categories according to their putative functions (Fig 9). A description of each category is provided in Supplementary Table S4. However, across all three species, a significant part of the DE baryzoite proteins were uncharacterized proteins, meaning hypothetical proteins predicted on genomic data. This shows that large portions of these genomes still need to be annotated (Fig 9).

**Figure 9.**
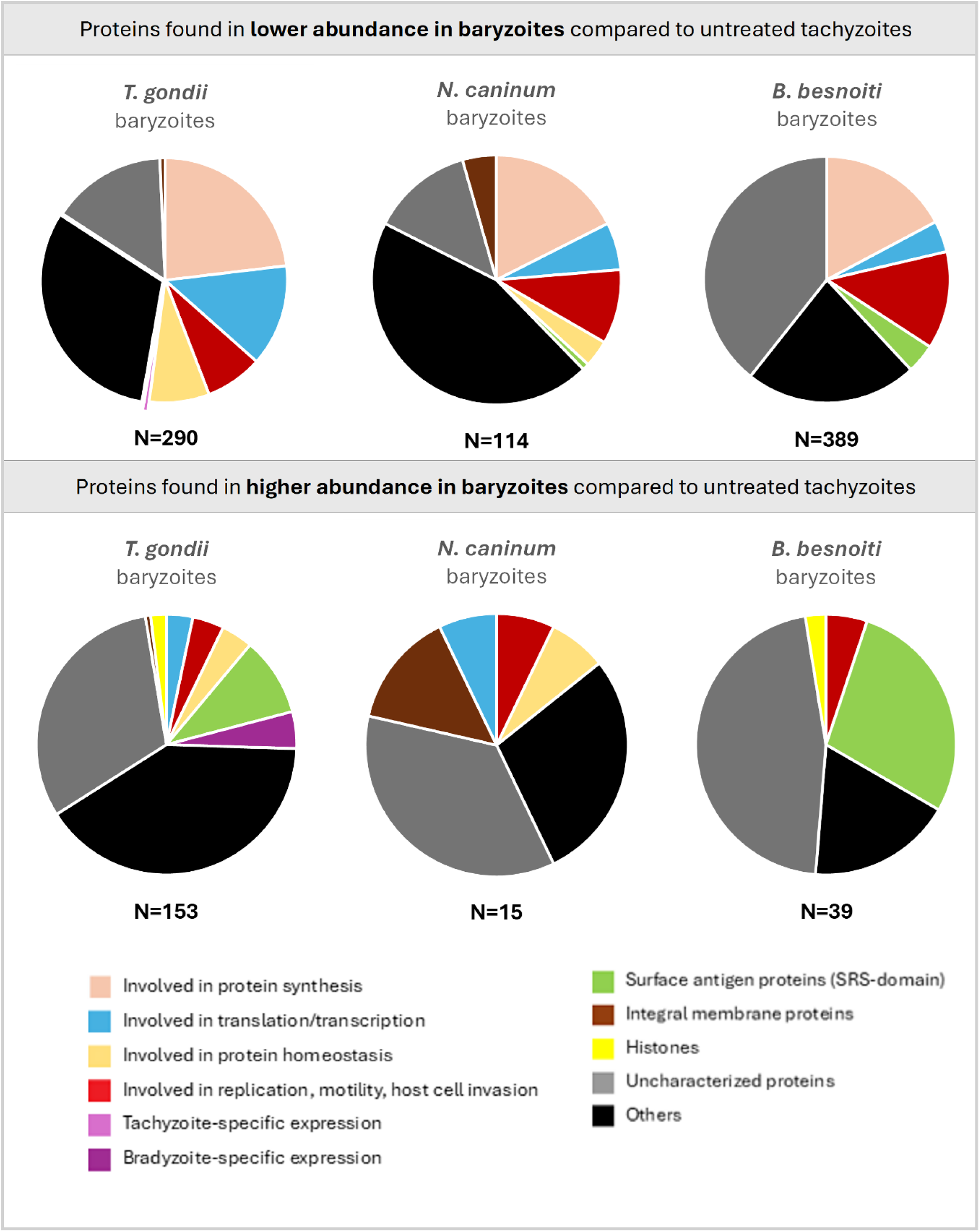
Differentially regulated proteins detected in baryzoites compared to untreated tachyzoites of *Toxoplasma gondii*, *Neospora caninum*, and *Besnoitia besnoiti*, categorized by their biological functions. “Uncharacterized proteins” include hypothetical proteins, and “Others” refer to detected proteins with unknown functions or those not classified within the defined categories. N = Total number of represented proteins.

Nevertheless, based on putative functional activity, a consistent pattern of protein downregulation was observed in baryzoites of all three species, with the same general groups of proteins exhibiting decreased levels, despite some parasite-specific differences. Following treatment, proteins involved in translation, transcription, replication, motility, and host cell invasion were less abundant in all parasites, with ribosomal proteins being most prominently affected: of the sixteen proteins that were found to be commonly expressed at lower abundance, only calmodulin and profilin did not represent ribosomal constituents, suggesting an impact on protein synthesis (Table 3). Several SRS-domain surface antigen proteins were expressed at reduced abundance in *B. besnoiti*, while only one was similarly affected in *N. caninu*m, and none in *T. gondii*. The list of SRS-domain surface antigen proteins differentially regulated in baryzoites exposure is given in Table 4. Reduced expression in *T. gondii* baryzoites was also noted for the tachyzoite markers enolase 2 (ENO2; TGME49_268850), lactate dehydrogenase 1 (LDH1; TGME49_232350), and calcium dependent protein kinase 1 (CDPK1; TGME49_301440), which is one of the BKI-targets identified to date (Fig 10A). These markers were not affected in *N. caninum* or *B. besnoiti*. CDPK1 orthologues in *N. caninum* (BN1204_044860) and *B. besnoiti* (BESB_015950) were detected in lower abundance in baryzoites compared to untreated parasites, although the variation was not statistically significant.

**Figure 10.**
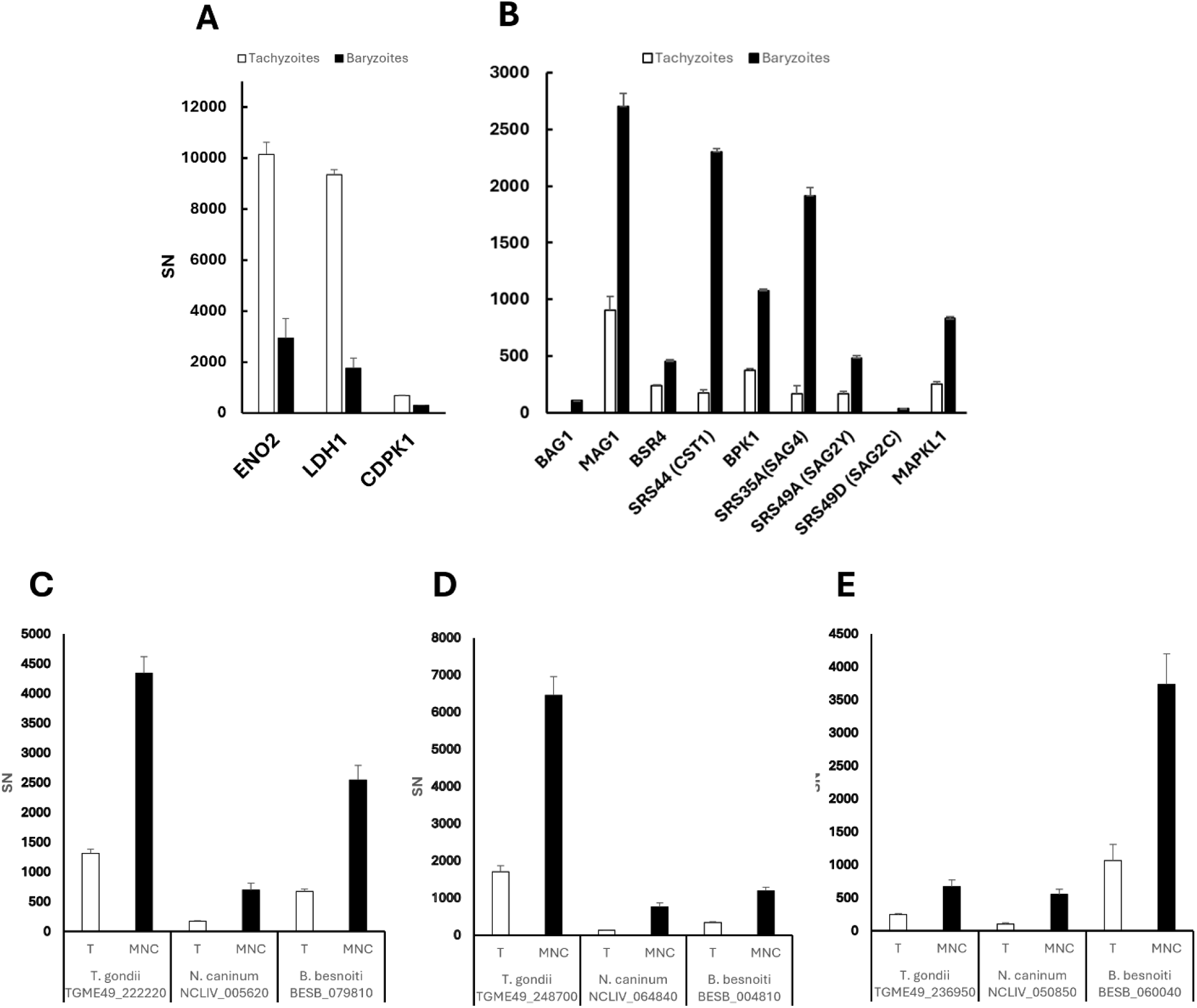
DE proteins found in lower (A) and higher (B–E) abundance in treated *T. gondii* compared to untreated cultures. DE proteins with significantly higher levels in treated cell cultures (baryzoites) compared to untreated cultures (tachyzoites) in *T. gondii*, *N. caninum*, and *B. besnoiti* include: (C) alveolin-domain intermediate filament IMC7; (D) alveolin-domain intermediate filament IMC12; and (E) an hypothetical protein. DE proteins were identified as described in Materials and Methods. Spectronaut intensities (SN) are given as mean ± standard deviation for three replicates. T – tachyzoites; MNC – multinucleated complexes or baryzoites.

**Table 3.** List of the sixteen proteins found at lower abundance in *T. gondii, N. caninum* and *B. besnoiti* baryzoites compared to untreated parasites (tachyzoites). Proteins are identified by ORF IDs listed in ToxoDB.

| Annotation in *N. caninum* | *N. caninum* | Orthologue<br>*T. gondii* | Orthologue<br>*B. besnoiti* |
| --- | --- | --- | --- |
| Profilin | NCLIV_000610 | TGME49_293690 | BESB_082700 |
| Calmodulin, putative | NCLIV_001300 | TGME49_305050 | BESB_025690 |
| Yml024wp-like protein, related | NCLIV_002780 | TGME49_207840 | BESB_022530 |
| 40S ribosomal protein S28, related | NCLIV_003680 | TGME49_209290 | BESB_023540 |
| 40S ribosomal protein S6 | NCLIV_012120 | TGME49_210690 | BESB_016090 |
| 40S ribosomal protein S15 | NCLIV_013320 | TGME49_213350 | BESB_046950 |
| 60S ribosomal protein L23a | NCLIV_015410 | TGME49_238010 | BESB_081300 |
| 40S ribosomal protein S7 | NCLIV_015880 | TGME49_239100 | BESB_082060 |
| 60S ribosomal protein L13 | NCLIV_024870 | TGME49_263050 | BESB_001210 |
| 30S ribosomal protein S12, related | NCLIV_030490 | TGME49_229670 | BESB_083100 |
| 30S ribosomal protein S15P/S13e, related | NCLIV_036280 | TGME49_270380 | BESB_053520 |
| 60S ribosomal protein L32, related | NCLIV_038780 | TGME49_267400 | BESB_050430 |
| 40S ribosomal protein S24 | NCLIV_052380 | TGME49_215460 | BESB_034520 |
| Ribosomal protein | NCLIV_052390 | TGME49_215470 | BESB_034510 |
| 60S ribosomal protein L6 | NCLIV_056680 | TGME49_313390 | BESB_074380 |
| 60S acidic ribosomal protein P0 | NCLIV_061830 | TGME49_218410 | BESB_039290 |

**Table 4.**
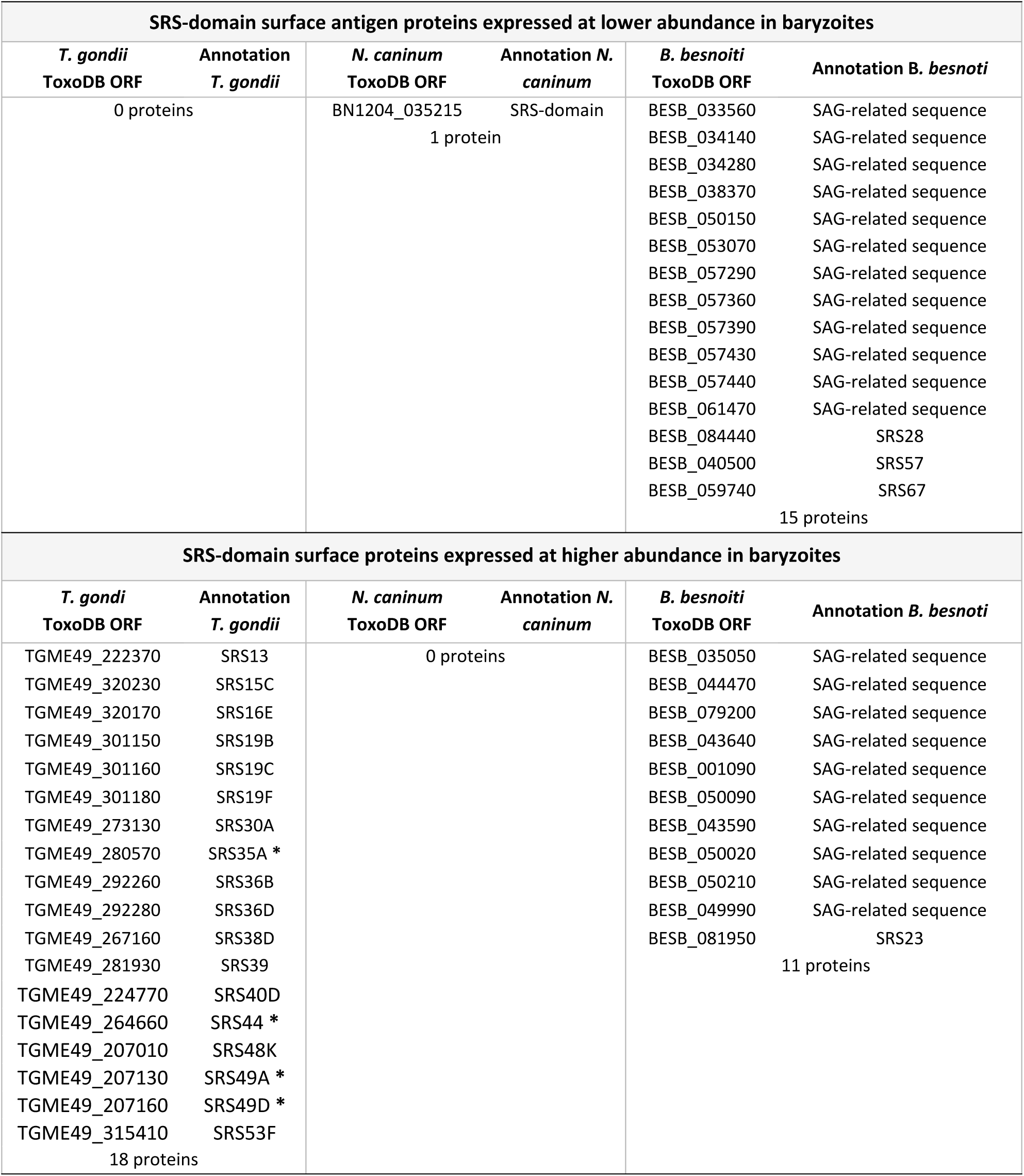
SRS-domain surface proteins expressed at low and high abundance in *T. gondii*, *N. caninum* and *B. besnoiti* baryzoites. ORF IDs correspond to those listed in ToxoDB. Bradyzoite-specific SRS proteins are marked with an asterisk (*).

Among the 153, 15 and 39 proteins that were detected at higher abundance in baryzoites of *T. gondii*, *N. caninum* and *B. besnoiti*, respectively, only three were upregulated in all three species. These include orthologues of the two alveolin domain containing intermediate filament proteins IMC12 (NCLIV_064840, BESB_004810, TGME49_248700) and IMC7 (NCLIV_005620, BESB_079810, TGME49_222220), and a hypothetical protein (NCLIV_050850, BESB_060040, TGME49_236950) (Figs 10C-E).

The 10 most highly abundant proteins of the 153 with increased expression levels in *T. gondii* baryzoites are shown in Table 5. In response to treatment, *Toxoplasma gondii* exhibits significant upregulation of specific histone variants, notably histone H3.3 (TGME49_218260), H2A.Z (TGME49_300200) and CENH3 (TGME49_225410) (Table 5). These variants play important roles in chromatin dynamics and gene regulation: H3.3 is associated with active transcription and chromatin remodeling, H2A.Z is implicated in transcriptional regulation and chromatin stability; and CENH3, a centromeric histone variant, is crucial for proper chromosome segregation during cell division (47). The increased levels of these histone variants and alveolin-domain intermediate filaments IMC7 and IMC12 in baryzoites suggest that BKI-1708 treatment induces specific coordinated changes in chromatin and cytoskeletal remodeling events, consistent with the observed alterations in cell cycle dynamics (Figs 2-6). Within the calmodulin-like protein family, the essential light chain ELC2 (TGME49_305050) (ortholog of *N. caninum* NCLIV_001300 and *B. besnoiti* BESB_025690) was downregulated across species (Table 3), whereas calmodulin (TGME49_249240) was specifically upregulated in *T. gondii*, possibly suggesting recruitment of an alternative calmodulin to support calcium signaling and stress adaptation (48). Among other proteins with increased abundance were bradyzoite markers such as TgBAG1 (TGME49_259020), the matrix antigen 1 TgMAG1 (TGME49_270240), as well as bradyzoite-specific SRS-domain proteins such as BSR4 (TGME49_320180), SRS44 (TGME49_264660), SRS35A (TGME49_280570), SRS49A (TGME49_207130), SRS49D (TGME49_207160), and the pseudokinase BPK1 (TGME49_253330), which mediates infectivity of *T. gondii* tissue cysts (Fig 10B). Moreover, BKI-1708 treatment resulted in increased expression of the secondary BKI-target MAPKL1 (TGME49_312570) (Fig 10B). The corresponding orthologs of TGME49_312570 in *N*. *caninum* and *B*. *besnoiti* are NCLIV_056080 and BESB_073700, respectively. Comparative analysis of the amino acid sequences of ATP binding sites suggests that they are highly conserved (Suppl. Fig 1). However, in our analysis, we could not detect the MAPKL1 orthologues NCLIV_056080 and BESB_073700 in the *N. caninum* and *B. besnoiti* proteomes, neither in tachyzoites nor in baryzoites.

**Table 5.** List of the ten most abundant proteins detected in *Toxoplasma gondii* baryzoites. Proteins are ranked according to the arithmetic mean of triplicates relative abundance (rAbu), with values presented as mean ± standard deviation (SD).

| ToxoDB ORF | Annotation | rAbu | SD |
| --- | --- | --- | --- |
| TGME49_218260 | histone H3.3 | 34537 | ± 6394 |
| TGME49_300200 | histone H2AZ | 25146 | ± 5105 |
| TGME49_225410 | histone H3 centromeric CENH3 | 13850 | ± 2712 |
| TGME49_249240 | calmodulin, putative | 4685 | ± 1467 |
| TGME49_248700 | alveolin domain containing intermediate filament IMC12 | 3150 | ± 503 |
| TGME49_222220 | alveolin domain containing intermediate filament IMC7 | 2556 | ± 361 |
| TGME49_226570 | hypothetical protein | 2417 | ± 399 |
| TGME49_270240 | MAG1 | 2334 | ± 874 |
| TGME49_280570 | SAG-related sequence SRS35A | 1506 | ± 636 |
| TGME49_236950 | hypothetical protein | 1261 | ± 337 |

No bradyzoite-specific markers were found to be expressed at higher levels in *N. caninum* or *B. besnoiti* baryzoites. In addition, two uncharacterized proteins (TGME49_226570 and TGME49_248360) were found at higher levels in treated *T. gondii* and *N. caninum*, but not in *B. besnoiti*. Their respective orthologues in *N. caninum* are NCLIV_046410 and NCLIV_064330.

## Discussion

Here, we present a comparative study of *T. gondii*, *N. caninum*, and *B. besnoiti* baryzoites. Baryzoites are multinucleated complexes that are formed upon exposure of these parasites to the bumped kinase inhibitor BKI-1708, a compound that had earlier shown promising efficacy against *N. caninum* and *T. gondii* infection *in vitro* and in pregnant mouse models (29). We thus report on the similarities and differences between the three species by focusing on the impact of BKI-1708 on ultrastructure, parasitostatic activity, the expression and localization of bradyzoite and tachyzoite markers, and proteomics.

Comparative TEM analysis showed that *in vitro* treatments of 2.5 μM BKI-1708 induced baryzoite formation in all three species, virtually no individual tachyzoites were found, showing that the changes observed concerned the entire population of parasites in a given culture. Inspection by TEM revealed many ultrastructural similarities between the three species. Overall, baryzoites represent a complex of newly formed, defective zoites that failed to complete cytokinesis. Consequently, disjunction and the formation of infective tachyzoites were inhibited. Parasites remained trapped within the host cell, while DNA-replication and nuclear division appeared to continue. In the case of BKI-1708, this resulted in the accumulation of nuclei that were typically clustered together to one side or to the periphery of a baryzoite, while the cytoplasm of newly formed zoites, which were separated from each other only by a single membrane, occupied the residual space. Thus, nascent apical complexes, conoids, and respective secretory organelles commonly found in apicomplexans, including rhoptries, dense granules and micronemes, were visible within the baryzoite matrix. In several instances, newly formed anterior ends of the parasites pointed outwards to the periphery. These cytoplasmic elements were most clearly discernible in *T. gondii* baryzoites. Overall, BKI-1708-induced baryzoites were structurally similar to those previously observed following treatments of *T. gondii* and *N. caninum* with the PP compound BKI-1294 (21, 28) and the AC compound BKI-1748 (18), and to those of *B. besnoiti* treated with a series of other PP and AC BKIs (49). Despite considerable changes induced by BKI-1708, baryzoites appeared to be viable and metabolically active, as indicated by the presence of mitochondria with a more or less intact mitochondrial matrix and cristae.

Viability of baryzoites was confirmed by treatment-and re-growth experiments, achieving similar results as previously obtained with *N. caninum* baryzoites induced by BKI-1294 (28). In that study, zoites lacking the major tachyzoite antigen NcSAG1 emerged from the periphery of these complexes after 10 days of drug removal, invaded neighboring cells and formed infective and proliferating tachyzoites. Under longer-term continuous BKI-1708 drug pressure over 30 days, *T. gondii* baryzoites remained morphologically intact and treatments did not result in clearance of infection. It is unclear, however, whether these forms were still metabolically active, and re-growth after drug removal was not assessed. Earlier studies had reported that viable parasites resumed proliferation upon drug removal after 20 days of treatment with 5 μM BKI-1294 (21), also suggesting ongoing metabolic activity for extended periods of time.

Despite the fact that all three species formed intracellular MNCs after BKI-1708 treatment, TEM revealed several distinguishing structural features in each organism. On one hand, BKI-1708 treatments resulted in a higher number of cytoplasmic vacuoles and lipid droplets in the *Neospora* and *Besnoitia* baryzoite cytoplasm compared to *T. gondii*. Cytoplasmic vacuolization and the formation of lipid droplets could be indicators of metabolic stress (4, 17, 50). Increased vacuolization suggests that BKI-1708 treatment had a more significant metabolic effect on *Neospora* and *Besnoitia* baryzoites compared to *T. gondii* baryzoites. Among the proteins identified in higher abundance in *N. caninum* and *B. besnoiti* baryzoites, several have potential roles in intermediary metabolism and apoptosis. In *N. caninum*, higher-abundance baryzoite proteins include a metallophosphoesterase (NCLIV_009030) and a 2OG-Fe(II) oxygenase (NCLIV_042790), enzymes implicated in various metabolic pathways in other organisms (51, 52). Additionally, reflecting on the roles of similar proteases in *T. gondii*, *Plasmodium falciparum*, and other eukaryotes, the cathepsin B-like protease (NCLIV_069550), found at higher levels in *N. caninum* baryzoites, likely contributes to lysosomal protein turnover, autophagy, and apoptosis regulation [39, 40]. In *B. besnoiti*, five proteins found at higher abundance in treated parasites could be linked to intermediate metabolism: i) BESB_026010 and BESB_025960, which encode the ribonucleoside-diphosphate reductase large chain, are involved in DNA synthesis and nucleotide metabolism, ii) indole-3-glycerol phosphate synthase domain-containing protein (BESB_033410), containing an indole-3-glycerol phosphate synthase domain, is suggested to promote essential amino acid biosynthesis, iii) melibiase subfamily protein (BESB_071320), a glycoside hydrolase that may facilitate carbohydrate turnover and iv) GDA1/CD39 (Nucleoside phosphatase) family protein (BESB_027400), a nucleoside phosphatase expected to influence nucleotide homeostasis and energy balance. Such changes in protein abundance could potentially explain/reflect the severe metabolic stress phenotype observed specifically in BKI-1708–treated *Neospora* and *Besnoitia*.

A second difference concerned the dense granules, which were much more numerous and smaller in size in *Neospora* and *Besnoitia* baryzoites. However, no alterations were noted in the expression levels of dense granule proteins in these parasites. The most notable difference, however, concerned the presence of an electron dense cyst wall-like structure that appeared at the periphery of the parasitophorous vacuole of *T. gondii* baryzoites after 6 days of BKI-1708 treatment. No such structures were visible after 4 days of treatment, suggesting that the material accumulated at the periphery in a time-dependent manner. In contrast, *N. caninum* and *B. besnoiti* baryzoites did not form a cyst wall. The build-up of a cyst wall by *T. gondii* baryzoites was confirmed by fluorescent labeling with the tissue cyst markers DBA and mAbCC2. The lectin DBA binds to the cyst wall glycoprotein 1 (CST1), a 116 kDa glycoprotein that plays a crucial role in maintaining cyst wall integrity and is essential for bradyzoite persistence within host tissues (55). The mAbCC2 recognizes a 40 kDa dense granule antigen in tachyzoites and recognizes a 115 kDa cyst wall protein that is upregulated during *T. gondii* bradyzoite differentiation (56, 57). In *N. caninum* infected host cells treated with sodium nitroprusside, both DBA and mAbCC2 had also labelled the walls of *in vitro* generated tissue cysts containing bradyzoites (46, 58). Furthermore, increased staining with mAbCC2 had also been observed in both *T. gondii* and *N. caninum* baryzoites induced by BKI-1294 treatment, with preferential peripheral staining on *T. gondii* and cytoplasmic staining in *N. caninum* baryzoites (21). However, upon treatment with BKI-1708, these antibodies failed to react with *N. caninum* or *B. besnoiti* baryzoites. Another bradyzoite marker, the bradyzoite antigen 1 (BAG1), a heat shock protein 30 (HSP30) related protein associated with tachyzoite-to-bradyzoite conversion (46, 58–60), was found to be expressed in *T. gondii*, but not in *N. caninum* or *B. besnoiti* baryzoites. We also performed immunofluorescence staining with antibodies against the major tachyzoite surface antigen 1 (SAG1), a tachyzoite specific antigen which plays an important role in host cell recognition, attachment, invasion, and modulation of the host immune response (61, 62). TgSAG1 labeling on the baryzoite surface in *T. gondii* got progressively weaker during BKI-1708 treatment and was completely diminished after 9 days. In contrast, in *N. caninum* baryzoites undergoing the same BKI pressure, the NcSAG1 signal persisted and remained visible even after 9 days of BKI exposure. Although orthologs of SAG1 and BAG1 have been identified in *B. besnoiti* (3, 63), antibodies reacted neither with *B. besnoiti* tachyzoites nor baryzoites. Overall, this indicated that the BKI-1708 induced baryzoites of *T. gondii* were more closely related to the tissue cysts generated by bradyzoites than their counterparts in *N. caninum* and *B. besnoiti*, which is in line with the results on the respective proteomes.

To obtain a more comprehensive overview on the proteins expressed in baryzoites of the three species; *T. gondii, N. caninum* and *B. besnoiti* infected HFF were treated with 2,5 μM of BKI-1708 for 5 days to induce the formation of baryzoites. Protein abundances were assessed by label-free quantitative proteomics using the Spectronaut algorithm (64) and analyzed in comparison to untreated tachyzoites. Proteins showing significant abundance changes compared to untreated samples were defined by a log₂ fold change ≥1 and p-value <0.05. The total number of proteins detected was highest in *B. besnoiti* (2755), followed by *T. gondii* (2670), and lowest in *N. caninum* (1461). A higher proportion of downregulated proteins was found in all treated strains, particularly in *B. besnoiti* (14.1 %) while the proportion of upregulated proteins in response to BKI-1708 treatment was rather low across all three species, ranging from 1 – 5 %. As mentioned above, the ultrastructural changes in *B. besnoiti* baryzoites, such as extensive vacuolization, partial mitochondrial alterations and abundant vesicular and electron dense material in the parasitophorous vacuole (Fig 4), were more pronounced than those observed in *T. gondii* or *N. caninum* (Figs 2 and 3). Consistently, proteomic profiling revealing that *B. besnoiti* exhibited i) the strongest downregulatory response and ii) highest upregulation of proteins potentially associated with metabolism and apoptosis under BKI-1708 treatment, suggesting a more pronounced shutdown of cellular functions compared to the other species. Nineteen orthologs showed consistent abundance changes across all three species, of which 16 were found at lower, and 3 at higher abundance.

Grouping DE proteins into functional categories (see Table S4) illustrated that proteins involved in the assembly and function of the ribosomes were most notably affected. Despite some parasite-specific differences, proteins involved in translation, transcription, replication, motility, and host cell invasion also exhibited lower abundance in all parasites, likely reflecting a common metabolic shutdown response associated with BKI-1708 treatments. Notable differences were detected with respect to the expression of SAG1-related sequence (SRS) proteins, with several SRS proteins less abundant in *B. besnoiti* baryzoites, only one SRS protein with decreased expression in *N. caninum* baryzoites, and no SRS domain-containing protein being downregulated in *T. gondii* baryzoites. This was not in agreement with the immunofluorescence results suggesting a progressive decrease of expression with time, although most effectively after 9 days of treatment. Thus, the discrepancy can be explained by the earlier timepoint of sample collection for proteomics. However, a substantial portion of proteins upregulated in response to BKI-1708 treatments were SRS proteins, namely 10 % in *T. gondii* and 28 % in *B. besnoiti* baryzoites. In contrast, no increased expression of SRS proteins was noted for the *N. caninum* baryzoite proteome. Earlier findings on the effects of BKI-1294 treatments of *N. caninum* had shown a different picture: of the 12 proteins exhibiting higher abundance in baryzoites, 6 were SRS domain containing proteins (27). Nonetheless, the bradyzoite-specific protein bradyzoite surface receptor 4 (BSR4) was detected in *N. caninum* baryzoites and not detected in untreated parasites, though it was not considered significantly differentially expressed according to the parameters of the applied DE test. Although variations in experimental settings between studies (e.g. incubation times, BKIs with different scaffolds, algorithms used for proteomics) prevent direct comparisons, other similarities in parasite responses were observed, such as reduced levels of housekeeping and intermediate metabolism proteins in BKI-1294 induced *N. caninum* baryzoites that align with observations in BKI-1708 induced baryzoites in *T. gondii*, *N. caninum* and *B. besnoiti* (27).

While immunofluorescence already indicated that *T. gondii* baryzoites, in contrast to *N. caninum* and *B. besnoiti,* move towards a bradyzoite-like expression pattern, this was confirmed on the proteomic level: tachyzoite markers such as enolase 2 (ENO2), lactate dehydrogenase 1 (LDH1) were downregulated. The same was observed for the BKI target CDPK1. Similarly, reduced expression of CDPK1 was also shown by immunofluorescence in *N. caninum* baryzoites treated with BKI-1294 (28). CDPK1 is essential for signaling leading to microneme secretion and thus host cell invasion, thus downregulation of its expression in baryzoites would have no impact on viability. In this study, no changes in expression levels of CDPK1 orthologues were found in *N. caninum* and *B.besnoiti* baryzoites. However, it is important to note that functional divergence and independent evolution of each species’ orthologs may influence their response levels under specific conditions.

Bradyzoite markers such as BAG1, matrix antigen protein 1 (MAG1), and several SRS proteins were expressed at higher levels in baryzoites. In addition, MAPKL1 (TGME49_312570), more recently identified as an alternative BKI-target in *T. gondii* (24) was more abundantly expressed in baryzoites than in tachyzoites. MAPKL1 is a coccidian-specific kinase that is localized to the pericentrosomal material and is essential for the regulation of centrosome duplication during endodyogeny. The presence of BKI-1708 most likely impacts on the activity of this kinase, which could be compensated for by upregulated expression in baryzoites, thus maintaining continued replication and nuclear division. Comparative sequence analysis for the corresponding orthologs of TGME49_312570, NCLIV_056080 in *N. caninum* and BESB_073700 in *B. besnoiti*, suggests that they are all MAPK type enzymes that are highly conserved. This would imply that *N*. *caninum* and *B*. *besnoiti* baryzoites should also have increased expression of NCLIV_056080 and BESB_073700 after BKI-1708 treatment. MAPKL1 orthologues in *B. besnoiti* and *N. caninum,* however, were not detected in the respective proteomes, neither in tachyzoites nor in baryzoites - suggesting that they are expressed at very low levels, or the protein is not stable under the conditions used. Also, noteworthy, serine palmitoyltransferase SPT2 (TGME49_290970), a key enzyme in ceramide biosynthesis, was detected in *T. gondii* baryzoites but not in untreated tachyzoites. It has been shown that TgSPT2 primarily produces long-chain dihydroceramides (dhCers), and deletion of TGME49_290970 does not notably impact on parasite growth or invasion *in vitro*. However, its presence becomes critical during chronic infection, where the absence of TgSPT2 results in reduced cyst formation in the brain, suggesting a role in parasite persistence and establishment of the intracellular niche (65). The absence of bradyzoite markers in *N. caninum* baryzoites may be attributed to the sensitivity limitations of the proteomic methods employed (66). Alternatively, it could reflect species-specific differences in bradyzoite formation. *N. caninum* bradyzoite formation *in vitro* is inherently more difficult to achieve, whereas certain *T. gondii* strains can even differentiate spontaneously, and bradyzoite formation is a well-characterized process (67, 68).

Among all identified proteins, 19 are commonly differentially regulated across all BKI-1708 induced baryzoites. Of these, 16 proteins are consistently detected at lower levels in baryzoites, 14 of which are ribosomal proteins, thus reflecting a general decrease in protein synthesis and a transition from an actively replicating state to a dormant form. In *N. caninum,* differential affinity chromatography using BKI-1748, a related AC compound, resulted in the identification of a number of BKI-1748-binding proteins involved in the interaction and modification of RNA, in particular, splicing, as well as proteins involved in DNA binding or modification and key steps of intermediate metabolism (69). Whether any of these proteins, or even nucleic acids, could act as alternative targets for other AC compounds such as BKI-1708 needs to be investigated.

Two additional commonly downregulated proteins were calmodulin (NCLIV_001300, TGME49_305050, BESB_025690) and profilin (NCLIV_000610, TGME49_293690, BESB_082700). Calmodulin is a small, Ca^2+^-binding protein. After binding to Ca^2+^, activated calmodulin can interact with various targets including motor proteins, ion channels, kinases, phosphatases, and membrane transport proteins, which apicomplexan parasites rely on for essential functions such as protein secretion, motility, host cell invasion and egress (70, 71). Of note, CDPK1, the kinase that is targeted by BKI-1708, contains a C-terminal Ca^2+^ calmodulin domain regulating its activity. In the presence of calcium, this domain undergoes a conformational change, allowing CDPK1 to activate and regulate downstream cellular processes including calcium-triggered microneme secretion (72, 73). Profilin is a small actin-binding protein localized at the apical end of tachyzoites, and is essential for gliding motility, host cell invasion, and active egress from host cells in *Toxoplasma gondii* (74). Profilin regulates the formation and depolymerization of short actin filaments in the inner membrane of the apical complex, enabling the translocation of myosins and adhesins (75, 76). *T. gondii* profilin is efficiently recognized by the immune system through binding to TLR11 and TLR12 and activates murine dendritic cells and macrophages to release IL-12, which in turn leads to the production of IFN-γ via the MyD88 dependent pathway and induction of Th1 biased immunity. Thus, profilins have been incorporated into vaccine candidates against *T. gondii* (77) and *N. caninum* infection (78).

Furthermore, orthologs of 3 proteins were detected at higher abundance across all strains. These include the alveolin domain containing intermediate filament protein IMC7 (orthologues: TGME49_222220, NCLIV_005620, BESB_079810) the alveolin domain containing intermediate filament protein IMC12 (orthologues: TGME49_248700, NCLIV_064840, BESB_004810), and an uncharacterized protein (orthologues: TGME49_236950, NCLIV_050850, BESB_060040). The inner membrane complex (IMC) is a double-membrane complex composed of alveolin-family proteins and essential for maintaining cell shape, motility, and host cell invasion, important for parasite stability and intracellular replication (79). Both IMC7 and IMC12 are essential for the structural integrity of the IMC in *T. gondii*. Their roles are particularly critical during the formation of daughter cells, replication and overall fitness (79). IMC7 and IMC12 filaments are typically incorporated into the mature cytoskeleton during the G1 phase of the cell cycle, when cytokinesis is complete (80). The expression of IMC proteins is tightly regulated during the cell cycle (81). In the context of baryzoites, the accumulation of these IMC-complex filament proteins in baryzoites could be explained by BKI interference with regulatory factors, such as transcription factors or kinases and/or arrest or delay of the G1 phase.

Another notable finding was the consistent detection of high levels of the uncharacterized protein TGME49_236950 (orthologues: NCLIV_050850, BESB_060040) in BKI-1708 induced baryzoites, which expression was downregulated in an artemisinin-resistant *Toxoplasma* strain (82). This hypothetical protein is highly conserved in apicomplexans, yet its function remains unknown. The hypothetical protein TgME49_236950 shares similarities with redox activators of prokaryotic corrinoid proteins, cobalt-containing cofactors involved in methyl-transfer reactions (reviewed in (82)). In *T. gondii* ME49 artemisinin-resistant strains, TgME49_236950 is present in lower levels, which could be linked to a gain-of-function with respect to these compounds — either through direct activation of artemisinins or by indirectly repressing proteins that interfere with their metabolic activation (82). Interestingly, in BKI-1708–induced baryzoites, this protein is consistently found at higher abundance. The increased abundance of TgME49_236950, NCLIV_050850, and BESB_060040 in *T. gondii, N. caninum* and *B. besnoiti* baryzoites, respectively, could reflect a role in stress adaptation and metabolic remodeling during tachyzoite-baryzoite conversion since its similarity to redox activators suggests a potential function in managing oxidative stress or participating in drug metabolism. Alternatively, this upregulation could result from stage-specific expression or compensatory pathway activation triggered by drug-induced signaling changes. Further functional studies would be required to clarify whether these orthologues play an active role in drug response, stage conversion, or broader redox and metabolic processes.

Finally, while this study has identified a number of features of baryzoites across *B. besnoiti*, *T. gondii*, and *N. caninum*, it is important to note that a substantial proportion of the proteomes are comprised of uncharacterized proteins. These proteins, "hypothetical proteins" are predicted based on open reading frames (ORFs) identified through genomic sequencing but lack experimental validation of their expression or functional characterization. This represents a substantial hurdle for clarification of pathways involved in this parasitostatic response to treatments with BKI-1708. Whether any of the proteins expressed at lower or higher abundance in baryzoites are indeed the ones directly involved in baryzoite formation is not known. Possibly, additional effects could contribute to this phenotype in a more indirect manner, such post translational modifications of stably expressed proteins that activate or inactivate certain functional activities important for the completion of cytokinesis of newly formed zoites and/or egress of infective tachyzoites. However, characterizing the formation of this drug induced stage and investigating the molecular pathways involved could offer new insights into the biology and persistence of cyst-forming apicomplexans.

## Conclusion

The consistent formation of baryzoites across *T. gondii*, *N. caninum*, and *B. besnoiti* and other apicomplexans in response to BKI treatment points to a conserved and regulated stress response, although with species-specific differences. Baryzoites are defined as a distinct dormant drug-induced stage in cyst-forming parasites, representing an alternative form between tachyzoites and bradyzoites, triggered and maintained by drug pressure *in vitro*. They are characterized by the presence of numerous nuclei resulting from continuous division, with an accumulation of newly formed daughter zoites within the host cell due to impaired cytokinesis. This stage is artificially induced and reversible, presenting ambiguous features of tachyzoites and bradyzoites simultaneously. The existence of baryzoites *in vivo* is uncertain, since no specific marker has been identified to date. However, similar multinucleated forms have been described in *T. gondii* upon treatment with diclazuril, a triazinone derivative that is targeting enzymes such as components of the mitochondrial respiratory chain and dihydrofolate reductase (83). Other repurposed compounds, such as the dopamine receptor pinozode and the estrogen antagonist tamoxifen also caused the formation of multinucleated forms of tachyzoites *in vitro* (84). Thus, the formation of a heavily enlarged parasitic stage that ensures survival in response to increased drug concentrations during prolonged periods of time might be more common than previously anticipated. Baryzoites grow exceptionally large by undergoing multiple rounds of nuclear division without forming complete daughter cells - reminiscent of schizonts, as seen in other Apicomplexa such as *Plasmodium spp*. Although these stages of asexual reproduction in *Toxoplasma gondii* are traditionally called meronts, the multi-nuclear baryzoite-phenotype suggests a functional resemblance to schizonts. To what extent baryzoites resemble the merogony stage that occurs in the cat intestine, during which the parasite undergoes several rounds of DNA replication and mitosis without immediately dividing, needs to be further investigated.

## Acknowledgments

We acknowledge the generous gifts of antibodies directed against TgSAG1 and TgIMC1 from Dominique Soldati-Favre, University of Geneva, Switzerland, anti-NcSAG1 from Camilla Björkman, University of Uppsala, Sweden, mAbCC2 and DBA from Adrian Hehl, University of Zürich, Switzerland.

